# Initiation of fibronectin fibrillogenesis is an enzyme-dependent process

**DOI:** 10.1101/2022.11.16.516843

**Authors:** Shay Melamed, Shelly Zaffryar-Eilot, Elisabeth Nadjar-Boger, Rohtem Aviram, Huaning Zhao, Wesal Yaseen-Badarne, Rotem Kalev-Altman, Dalit Sela-Donenfeld, Oded Lewinson, Sophie Astrof, Peleg Hasson, Haguy Wolfenson

## Abstract

Fibronectin fibrillogenesis and mechanosensing both depend on integrin-mediated force transmission to the extracellular-matrix. However, force transmission is in itself dependent on fibrillogenesis, and fibronectin fibrils are found in soft embryos where high forces cannot be applied, suggesting that force cannot be the sole initiator of fibrillogenesis. Here we identify a nucleation step prior to force transmission, driven by fibronectin oxidation mediated by lysyl-oxidase enzyme family members. This oxidation induces fibronectin clustering that promotes early adhesion, alters cellular response to soft matrices, and enhances force transmission to the matrix. In contrast, absence of fibronectin oxidation abrogates fibrillogenesis, perturbs cell-matrix adhesion, and compromises mechanosensation. Moreover, fibronectin oxidation promotes cancer cells colony formation in soft agar as well as collective and single-cell migration. These results reveal a force-independent enzyme-dependent mechanism that initiates fibronectin fibrillogenesis, establishing a critical step in cell adhesion and mechanosensing.

## Introduction

Cell adhesion to the extracellular matrix (ECM) is a fundamental property that is essential for multiple cellular processes including migration, proliferation, and differentiation. As a result, cellular functions often depend on alterations within the ECM, including changes in its composition, structure, and mechanical properties^1, 2^. Fibronectin (FN) is an essential ECM protein that promotes cell adhesion to the matrix^3, 4^, and is also an important regulator of ECM alterations owing to its binding to numerous ECM proteins, including collagen^5^, fibrillin^6^, fibrinogen^7^, and Tenascin-C^8^. FN is secreted as an inactive dimer and it promotes matrix assembly and integrin-mediated cell adhesion to the matrix in its active multimeric fibrillar form^9, 10^. The dimeric to pro-fibrillar transition, which activates the initiation of FN fibrillogenesis, is considered to be driven by intracellular forces generated and transmitted via the actomyosin cytoskeleton to the extracellular FN molecules through transmembrane integrins^11–13^. When generated, the forces lead to conformational changes within FN, exposing cryptic self-association binding sites which allow adjacent dimers to interact, thus gradually promoting the assembly of FN fibrils^14, 15^. This process is associated with growth of nascent integrin adhesions into mature focal-and/or fibrillar-adhesions^16, 17^. Indeed, in situations where high forces cannot be generated, as in cultured cells grown on soft matrices, FN fibrillogenesis is inhibited, resulting in reduced cell adhesion^18^. In developing embryos, fibril assembly depends on cadherin-dependent enhanced contractility that leads to integrin-FN translocation^19^. However, in very early stages of embryonic development, the embryo rigidity is so low^20, 21^ such that it is not expected to provide sufficient resistance to support transmission of high cellular forces to the matrix. Thus, the observed FN fibrils at such early embryonic stages^22, 23^, which are essential for facilitating cell adhesion and migration, appear to depend on additional mechanisms which can overcome and compensate for the low rigidity and the resulting inability to exert force. Here we set to dissect this longstanding conundrum. We found a force-independent mechanism that is essential for the initiation of FN fibrillogenesis.

## Results

### Lysyl oxidases promote FN fibrillogenesis and cell spreading on soft matrices

The presence of FN fibrils in early, soft, embryos whose stiffness cannot support transmission of high forces that are involved in integrin-dependent FN conformational changes, suggests that an additional mechanism is involved in FN fibrillogenesis. Our previous finding that members of the Lysyl oxidase enzyme family bind FN, that LOX-Like 3 (LOXL3) oxidizes it, and that its enzymatic activity induces FN-dependent integrin activation^24^, raised the possibility of an enzyme-dependent initiation of fibrillogenesis, a force-independent mechanism which relies on enzymes and their substrates’ availability. In line with this notion, we found that in cultured mouse embryonic fibroblasts (MEF) expressing GFP-tagged FN^25^, FN fibrils are observed under β1-integrin adhesions and LoxL3 is localized at the cell periphery together with small FN patches (Figure 1A, arrowheads). Analysis of the spatial interactions^26^ between β1-integrin and LoxL3 demonstrates that the two proteins are colocalized specifically along the cells’ edges (Figure 1B).

**Figure 1.**
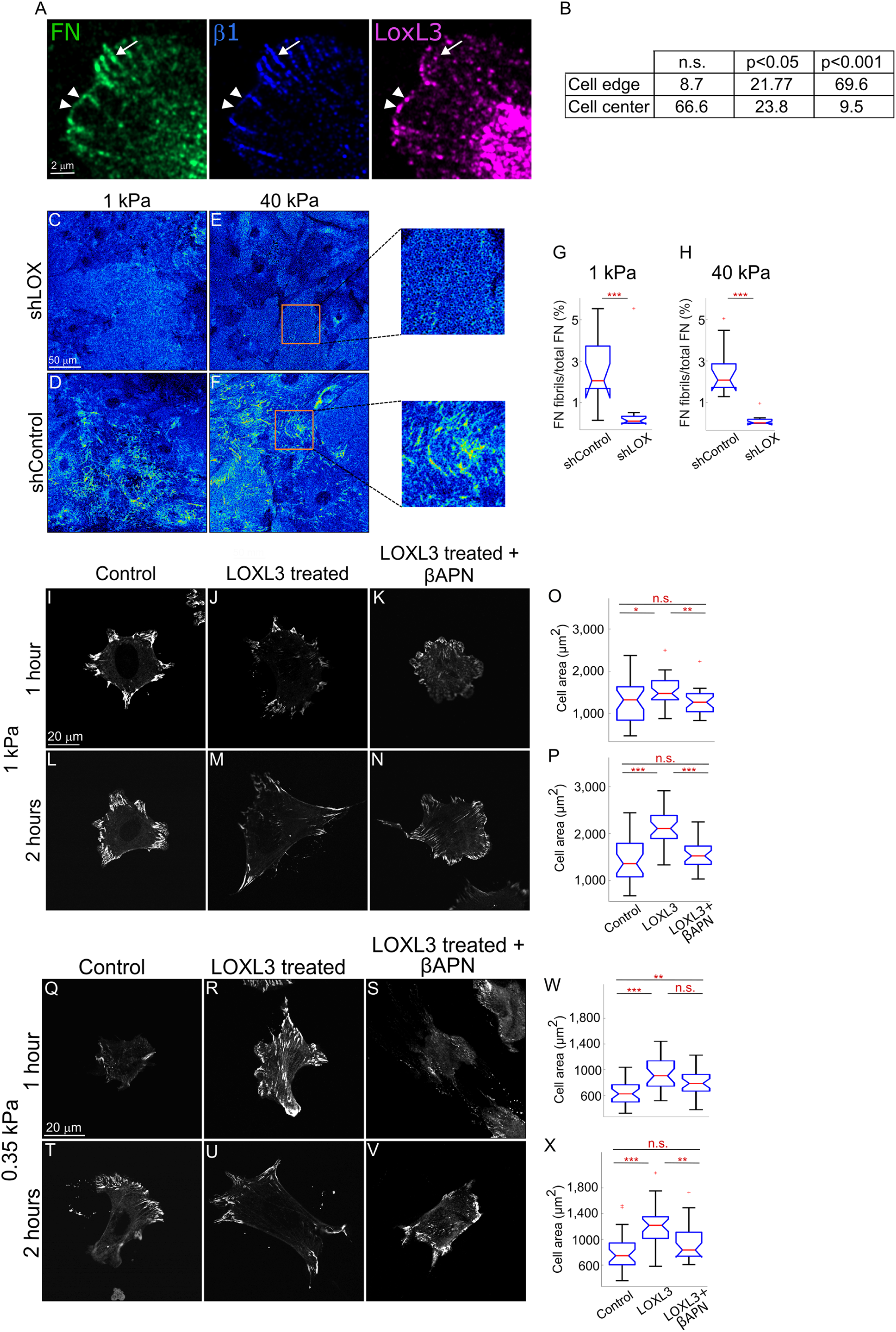
Lysyl oxidases regulate FN fibrillogenesis on soft matrices. MEF expressing FN-GFP stained for β1-integrin and LoxL3. FN-GFP clusters and LoxL3 clusters co-localize at the cell boundary (arrowheads) whereas fibrillar FN localizes under mature β1-integrin adhesions (arrows) (A). Summary of the percentage of cells for which co-localization between β1-integrin and LoxL3 was identified at the indicated significance levels, showing significantly higher co-localization at the cell edge versus the cell center (B). FN immunostaining, color-coded for intensity (blue – low, yellow – high), of *shLOX* and *shControl* HASMC on 1 kPa (C-D) and 40 kPa (E-F). Magnification of representative regions in E and F. Quantification of FN fibrils show significant reduction following LOX knockdown on both 1 kPa and 40 kPa (G-H, n>40). Pax-GFP on sham-and LOXL3-treated FN with or without βAPN, seeded on 1 kPa (I-N) or 0.25 kPa (Q-V) gels and fixed after 1 hour (I-K, Q-S) and 2 hours (L-N, T-V). Quantifications of cell area show significant differences between cells seeded on treated FN regardless of the rigidity (O-P, W-X, n>28). **p<0.0001, **p<0.001, *p<0.05, n.s. = non-significant.

Notably, LoxL3 is expressed in neural crest cells (Figure S1A) and following its deletion (*LoxL3^Δ/Δ^*) the mutant embryos display a cleft palate phenotype and reduced neural crest cell numbers at branchial arches (Figure S1B-H), both of which are FN-associated phenotypes^27, 28^. Further, abnormal FN-fibril formation along the somitic boundaries is also observed in *LoxL3^Δ/Δ^* mutant embryos (Figure S1I-K). Collectively, these findings suggest a role for Lysyl oxidases in FN fibrillogenesis.

To test the relationship between substrate rigidity, FN fibrillogenesis, and the Lysyl oxidases, we took advantage of primary human aortic smooth muscle cells (HASMCs), that a) normally reside on a rigid artery wall whose stiffness, depending on stretching, is 15-88 kPa^29^; and b) primarily express the lysyl oxidase family member, LOX^30^. Following infection with short hairpin RNA against *LOX* (*shLOX*) or scrambled RNA (*shControl*) (Figure S1L), we cultured the cells for 24 hours on 1 kPa or 40 kPa FN-coated surfaces and monitored FN-fibrils by immunostaining. Whereas *shControl* cells displayed significant FN fibril formation, cells devoid of LOX, despite secreting FN to the same extent (Figure S1M), did not form any FN fibrils on both rigidities (Figure 1C-H).

We next wished to carry out the converse experiment and test whether pre-treatment of FN with lysyl oxidases overrides rigidity sensing and promotes early cell adhesion on soft matrices. Towards that end, we used recombinant human LOXL3 or LOXL2, and treated FN-coated 1.0 and 0.35 kPa surfaces with either enzyme. MEFs stably expressing Paxillin-GFP (Pax-GFP) were then added onto the plates and fixed after 60 and 120 minutes on the sham-, LOXL3-or LOXL2-treated FN (Figure 1I-X and Figure S1N-ZC). FN fibrils were undetectable at these early times of cell spreading and therefore we monitored cell areas as readouts of cellular interaction with FN. This analysis revealed a significant increase in the areas of cells plated on the LOXL3-or LOXL2-treated FN in comparison to the sham-treated FN or to those treated with the enzyme mixed with the pan LOX inhibitor beta-aminopropionitrile (βAPN)^31^.

Previous work has noted significant differences in mechanosensing whether the ECM proteins were physically adsorbed or chemically cross-linked to the substrate as cell-generated forces can pull-off ECM molecules from the substrates in the former scenario^32^. Although we used soft silicone substrates (CY 52-276) on which this phenotype does not occur^33^, to verify that the above observations are not due to crosslinking of FN to the underlying gel via the LOX family members but rather due to a possible change in FN, we crosslinked LOXL3-treated or untreated FN to 0.35 kPa gels and monitored cell spreading after 2 hours. We found that in a similar manner to that observed above, also in this case, the treatment of FN with a LOX family member enhanced cell spreading compared to the untreated control (Figure S1ZD), demonstrating that the crosslinking step failed to significantly enhance the link between FN and the substrate beyond the physisorbed FN. Altogether, these results suggest a general role for the enzymatic activity of lysyl oxidases that promotes FN fibrillogenesis and cell adhesion.

### LOX-treated FN promotes nascent adhesion formation

The above results suggest that LOX family members modulate the cells’ response to the external rigidity by initiating FN fibrillogenesis and enhancing adhesion formation. We therefore wished to test whether the direct treatment of FN with a lysyl oxidase affects the mechanism by which cells perform mechanosensing of ECM rigidity. We postulated that the enzymatic activity of lysyl oxidases promotes a structural change in FN that allows creating more stable nascent adhesions through which early mechanosensing of matrix rigidity occurs^34, 35^. To test this, we plated cells on arrays of deformable PDMS pillars, with 0.5 µm diameter and 2.3 µm height (bending stiffness 1.5 pN/nm^36, 37^, equivalent to ∼2 kPa^38^), and performed live-cell time-lapse imaging of early attachment and spreading of the cells (Figure 2A-C). To test the hypothesis that LOX-induced local structural change in FN enhances nascent adhesion formation, we coated the pillars with FN at a density which was less than saturation level (1 µg/ml, equivalent to ∼25% coverage of the pillar tops). Analyses of early times of cell edge interactions with individual FN-coated pillars revealed repeated catch-and-release events which were three standard deviations above the mean noise level (Figure 2A). We analyzed the strength of the linkage in each of these events by measuring the pillar displacement level at the time of release, translating it to force. Comparison of the histograms of release forces between sham-and LOXL3-treated FN showed that the latter displayed significantly more events per unit time at 2 pN (Figure 2B), consistent with the formation of integrin-mediated FN-actin 2 pN bonds in the presence of trimers of FN’s integrin-binding domain FNIII7-10, but not with monomers^39^.

**Figure 2.**
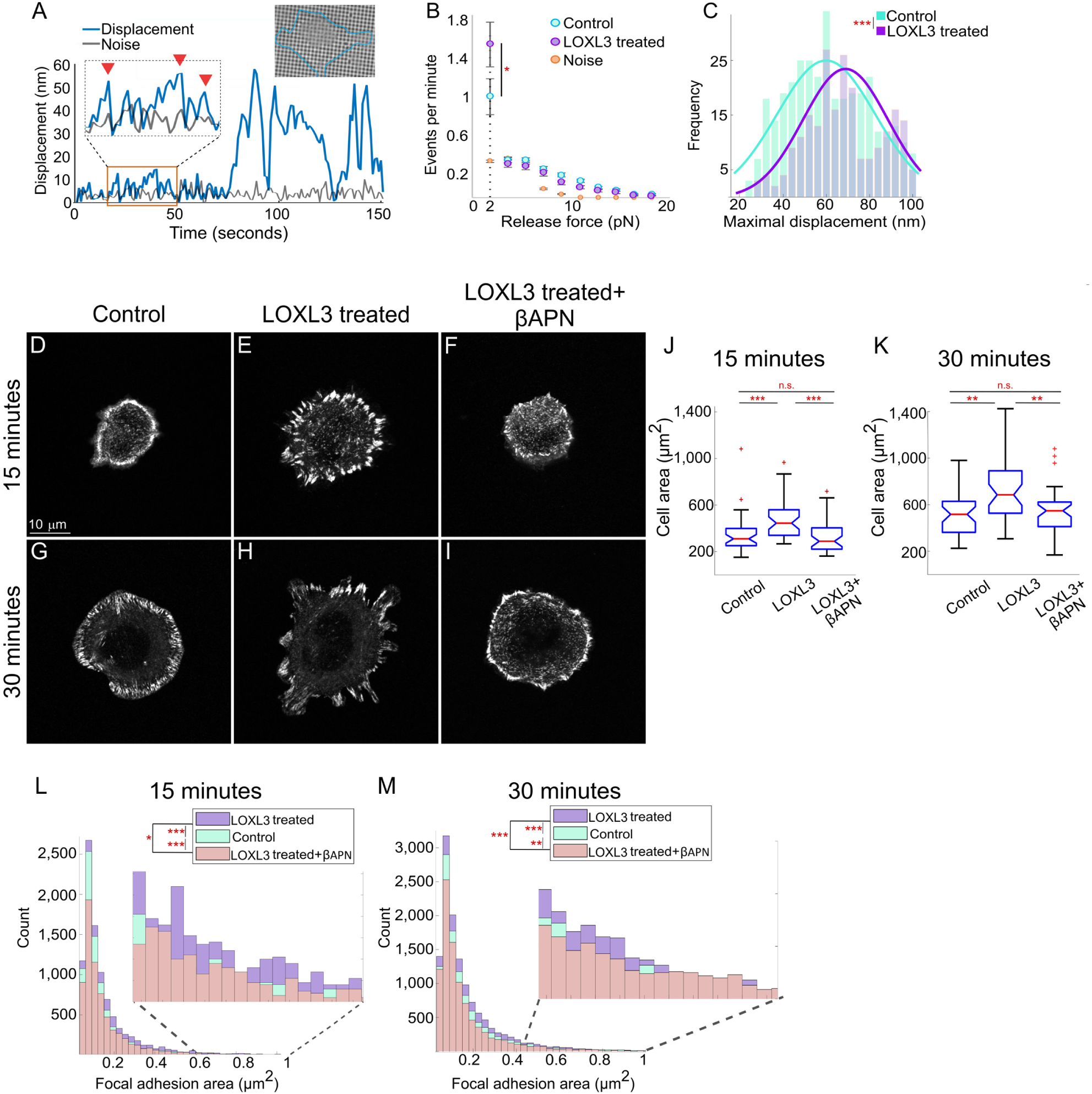
LOX family stimulate adhesion initiation. Representative image of a cell on pillars and a graph of pillar displacement; the red arrowheads point to early catch-and-release events (A). Histograms of the pillar release force during catch-and-release events by cells seeded on sham or LOXL3-treated FN, demonstrating enhanced formation of 2 pN bonds upon LOXL3 treatment (B, n>243). Histogram of the maximal displacement of pillars by cells seeded on LOXL3-treated FN (C, n>243). Pax-GFP on sham-and LOXL3-treated FN with or without βAPN, seeded on glass and fixed after 15 (D-F) and 30 minutes (G-I). Quantifications of cell area show significantly enhanced spreading of cells seeded on treated FN compared to controls (J-K, n>39). Histogram of focal adhesion area in cells seeded on sham-or LOXL3-treated FN demonstrate that LOXL3 treatment promotes formation of larger adhesions (L-M). ***p<0.0001, **p<0.001, *p<0.05, n.s. = non-significant.

These results are suggestive of formation of more closely packed FN clusters following LOXL3 treatment. Further, analyses of the longer-term pillar displacements that succeeded the initial catch-and-release events showed a significant shift toward higher values of the maximal displacement histogram in LOXL3-treated compared to control FN (Figure 2C). As control, we coated the pillars with laminin, an integrin ligand which interacts with members of the Lox family, yet which is not oxidized by them^24^. Under these conditions, the increments in the maximal pillar displacements were not observed (Figure S2A-B), highlighting the specificity of the oxidation process of FN. Together, these results suggest that the ability to form FN-dependent nascent adhesions on soft surfaces is enhanced upon LOX treatment, which assists in overcoming the low rigidity of the matrix (Figure 1).

We next considered the fact that nascent adhesion formation even on very stiff matrices requires at least four RGD (Arg-Gly-Asp) ligands that are localized within ∼60 nm from each other^40, 41^. Thus, if LOX treatment led to an adhesion promoting structural change in FN, one should expect to see a difference in early adhesion even on stiff surfaces such as glass whose stiffness is at the GPa range. Towards that end, following enzymatic treatment, FN was coated on glass coverslip and Pax-GFP cells were added and shortly after (15 or 30 minutes) fixed and analyzed. Quantifications of adhesion sizes at 15 minutes showed a significant increase in the relative count of adhesions larger than 0.1 µm^2^ on LOXL3-or LOXL2-treated FN and to a concomitant increase in cell spreading area (Figure 2D-F, J, L and Figure S2C-D, G, I).

Importantly, the increases in adhesion sizes and cell areas were blocked when βAPN was added to the FN-LOXL3 mix throughout the reaction (Figure 2I, J, L). A similar trend was observed at 30 minutes, albeit to a lesser extent (Figure 2G-I, K, M and Figure E-F, H, J), altogether suggesting that the primary effect of the enzymatic treatment occurred at the very early stages of adhesion assembly. A comparable effect was observed with two other fibroblast cells (human dermal fibroblasts and mouse embryonic fibroblasts; Figure S2K-R), indicating that the effect of lysyl oxidases on adhesion to FN was not specific to a particular cell line. To further verify the specificity of the LOX-FN adhesion effects, we carried out a similar experiment using LOXL3-treated laminin-coated plates. No change in cell area was observed following this treatment (Figure S2S-V), reinforcing the notion that the lysyl oxidases specifically modulate FN-dependent cell adhesion.

Notably, since α5β1 integrin can interact with LOX enzymes^42^, we wished to verify that the observed effect on adhesion occurred through changes in FN and not through direct integrin activation by residual enzyme that remained attached to FN (Figure S3A). Indeed, when βAPN was added following the FN treatment, no difference was observed in cell spreading (Figure S3B-F, J), demonstrating that the enzymatic activity of LOXL3 directly affects FN and that the presence of the enzyme by itself (when inactive) has no effect. Taken together, the similar results observed following LOXL2 and LOXL3 treatment (Figure 2 & Figures S2 and S3), and the observations on LOX effects on FN fibrillogenesis (Figure 1 & Figure S1), reinforce the idea of a general enzymatic role among LOX family members in affecting cell adhesion through initiation of FN fibrillogenesis.

### LOX induces FN clustering

We next wished to directly test whether alteration to FN organization occurs following LOX treatments. To this end, we deposited treated and untreated FN samples, as well as fresh FN, on glass coverslips at low density (100 ng/ml). After immunostaining the samples, we imaged regions completely devoid of cells using direct stochastic optical reconstruction microscopy (dSTORM). We then used particle analysis to identify FN clusters in the dSTORM images (Figure 3A-B). The histograms of the detected FN clusters indicate that LOXL3 treatment led to significant increase in the number of clusters larger than 0.009 µm^2^ compared to fresh or sham-treated FN (Figure 3C). Collectively, these results indicate that LOX treatment enhances FN clustering, which results in cells perceiving the matrix as stiffer.

**Figure 3.**
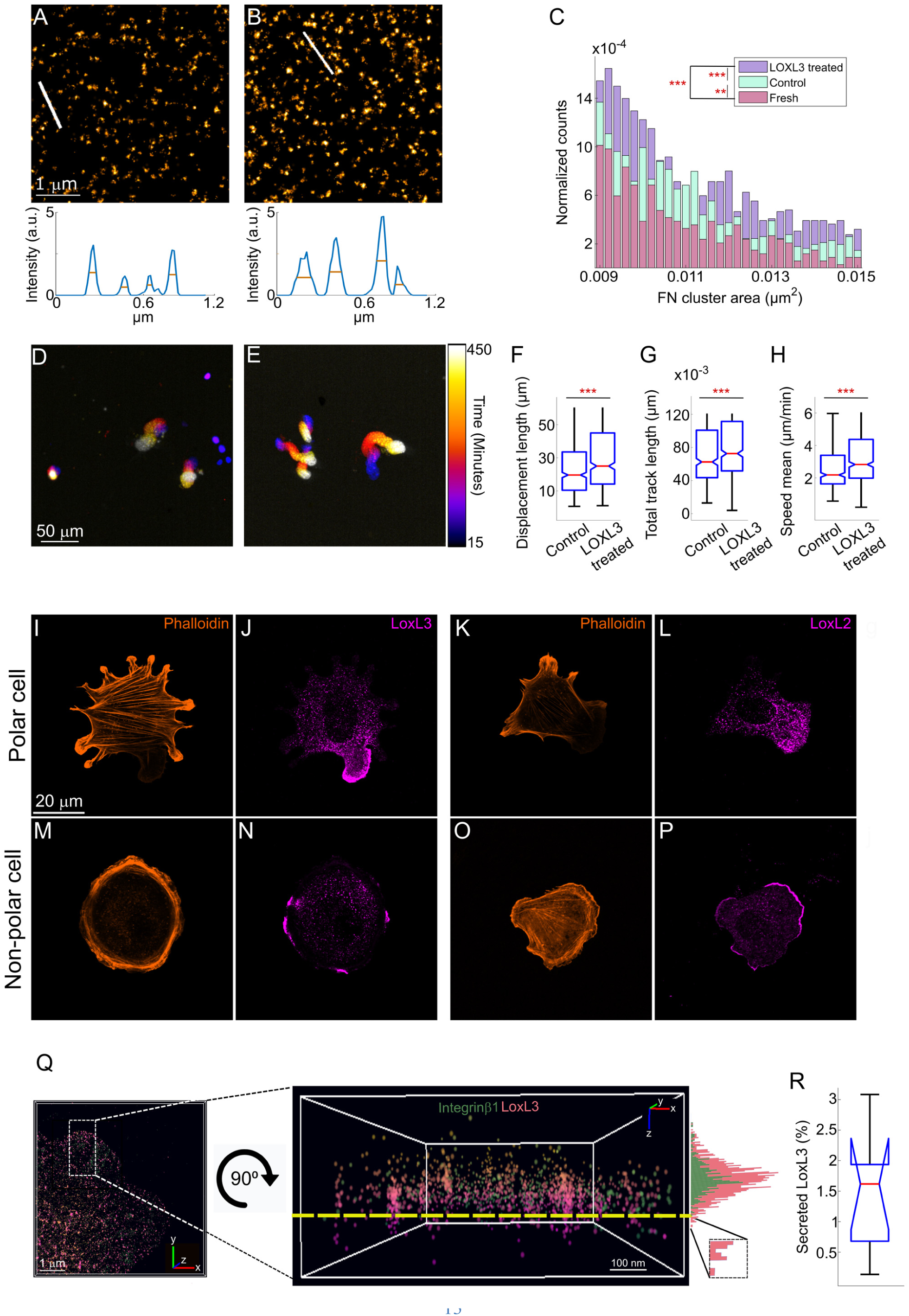
LOX induces FN clustering. Representative dSTORM images of sham-and LOXL3-treated FN nanodomains; the curves below are the intensity values along the white lines shown in each image (A-B). dSTORM analysis of FN cluster area shows a significant increase following LOXL3 treatment (C). Representative color-coded images generated from time-lapse videos tracking fluorescence-labeled nuclei on sham-and LOXL3-treated FN (D-E). Quantifications of migration parameters show significant increases in the measured parameters of cells seeded on LOXL3-treated FN (F-H, n>426). LoxL3 and LoxL2 immunostaining of non-polar cells (I-L) and polar cells (M-P) showing that the Lox family members are localized to lamellipodial regions. 3D dSTORM image and zoom-in x-y or x-z representation of the cell edge using β1-integrin (green) and LoxL3 (pink) staining. The histograms on the right represent the z-axis distribution of both proteins, showing the extension of LoxL3 closer to the coverslip, i.e. secretion of LoxL3 below the dashed yellow line (Q). Quantification of the percent of LoxL3 molecules localized below the cell leading edge (R, n=17). ***p<0.0001, **p<0.001.

The observation that treatment of FN with lysyl oxidases promotes FN clustering and induces cell adhesion raised the hypothesis that LOXL3-FN treatment leads to increased maturation and stabilization of nascent adhesions during early stages of cell interactions with the matrix. To test this hypothesis, we seeded Pax-GFP cells on LOXL3-treated and untreated FN and monitored nascent adhesion lifetimes using live-cell imaging. We find that both the rates (Figure S3G) and total increase in adhesion area (Figure S3H) are significantly increased following LOXL3 treatment.

Next, since adhesion stabilization plays a major role in maintenance of a leading edge in cell migration^43^, we postulated that the larger FN clusters following treatment would lead to improved migration. We therefore plated Pax-GFP cells on sham-and LOXL3-treated FN and performed live-cell imaging to track their migratory patterns and quantify the overall displacement length, migration distance, and speed (Figure 3D-H). All three parameters were significantly increased following LOXL3 FN-treatment, demonstrating that not only were the cells able to rapidly generate adhesions on the treated FN but importantly, that the cells’ migration pattern was more persistent (evident by the increased displacement length), further reinforcing the notion that the adhesions that form on the LOXL3-treated FN are more stable.

The overall results thus far suggest that exogenous FN treatment with a member of the lysyl oxidase family promotes a faster and more robust cell adhesion. As members of the family are secreted enzymes, we next wished to test whether their secretion is localized to cellular regions where such robust integrin-dependent adhesions are mostly required. To that end, Pax-GFP cells were immunostained for LoxL2 or LoxL3 following 60 minutes of spreading. We find that in polarized cells, LoxL2 and LoxL3 expression is typically localized to the most prominent lamellipodial protrusion of the cell (Figure 3I-L). Surprisingly, even in non-polarized cells, narrow stripes in lamellipodial regions were also observed (Figure 3M-P). Further, 3-dimensional (3D) dSTORM analysis demonstrates that these matrix modifying enzymes are specifically secreted from the leading edge (Figure 3Q-R). Taken together, these results support a model whereby localized secretion of LOX enzymes at the cell edge enhances FN clustering, thereby supporting nascent adhesion formation and persistent cell migration.

### FN oxidation is essential for FN fibrillogenesis

Since a) Lysyl oxidases’ enzymatic activity is required for FN-mediated cell adhesion and migration; b) treatment of FN with LOXL3 promotes its clustering; and c) LOXL3 oxidizes FN^24^, prompted us to further explore the oxidation and dissect its necessity. Liquid chromatography–mass spectrometry (LC-MS/MS) analysis identified three lysine residues (K116, K2391, K2401) that were specifically oxidized by both LOXL2 and LOXL3 (Figure 4A, Figure S4A-D and Tables S1-2). Since no active recombinant LOX enzyme is available, we cultured HASMC (which primarily express LOX, as mentioned above) and submitted the ECM they secrete to the LC-MS/MS analysis. We find that in a similar manner to the two LOX-like proteins, FN secreted by HASMC is also specifically oxidized at lysine 116 (Table S3), altogether demonstrating the specificity of the reaction and reinforcing the above observations of the shared FN-dependent activities the LOX enzymes carry. Analysis of the oxidized lysine residues demonstrates that they are all localized to FN type 1 (FN1) repeats (FN1 repeat #2 and FN repeat #12), one close to the FN N-terminus and the other in its C-terminal region. Both regions encompassing these lysine residues are highly conserved in vertebrate FN (Figure 4B) suggesting that they play an important role in FN activity.

**Figure 4.**
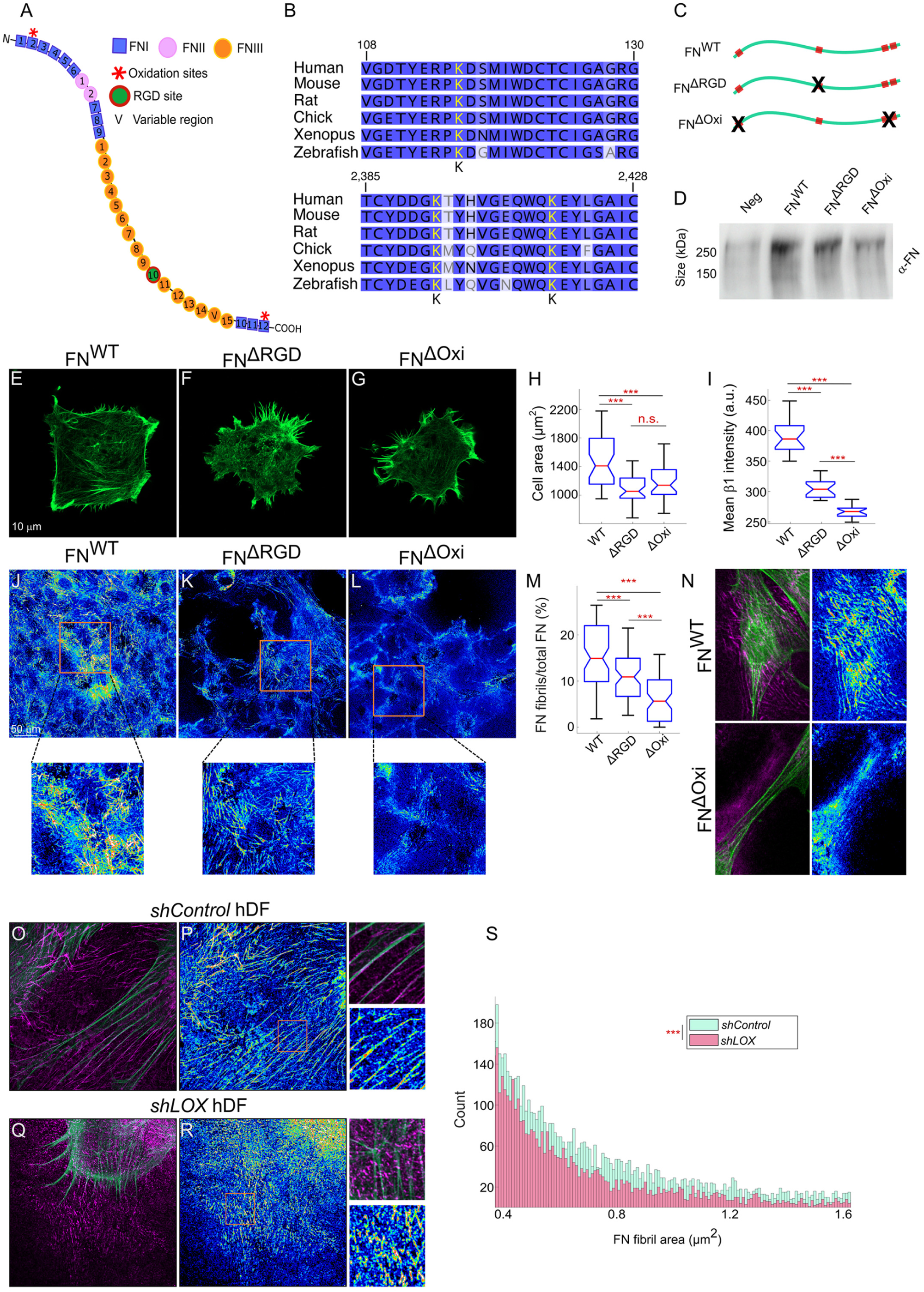
LOX is essential for FN fibrillogenesis. Schematic of a FN monomer with the oxidized lysine residues marked by red asterisks (A). Conservation of the three oxidized lysines among vertebrates (B). Schematic representing the two FN constructs that were generated (C). Western blot against FN from conditioned medium of cells expressing FN^wt^, FN^ΔRGD^ and FN^ΔOxi^ (D). Phalloidin staining of FN^wt^, FN^ΔRGD^ and FN^ΔOxi^ cells seeded on glass without external FN coating and fixed after 1 hour (E-G). Quantification of cell area shows a significant decrease in FN^ΔOxi^ cells (H, n>39). Quantification of mean fluorescence intensity of β1-integrin staining per cell shows a significant decrease in FN^ΔOxi^ cells (I, n>20). FN immunostaining, color-coded for intensity (blue – low, yellow – high), of cells expressing FN^wt^, FN^ΔRGD^ and FN^ΔOxi^ after 24 hours of seeding on glass without external FN coating; magnification of representative regions shown below (J-L). Quantification of FN fibrils show a significant decrease in FN^ΔOxi^ cells, revealing a critical contribution of LOX family to FN fibrillogenesis (M, n>71). Representative images of cells expressing FN^wt^ and FN^ΔOxi^ after 24 hours of seeding on glass without external FN coating, stained for F-actin and FN (color-coded for intensity), showing the lack of FN fibrils even when stress fibers are assembled in FN^ΔOxi^ cells (N). Actin and FN immunostaining (color-coded for intensity) of *shControl* (O-P) and *shLOX* (Q-R) Human dermal fibroblasts seeded on glass without external FN coating and fixed after 24 hours, demonstrating the inability of *shLOX* cells to form FN fibrils even when actin stress fibers are assembled. Images on the right are zoom-ins of the orange boxes in each image. Histogram of the area of FN fibrils generated by *shControl* or *shLOX* human dermal fibroblasts, demonstrating the significance of LOX in the long-term FN fibrillogenesis process (S). ***p<0.0001, n.s. = non-significant.

To directly test the role that these three lysine residues play in cell adhesion and FN fibrillogenesis, we generated a lentiviral construct expressing a mutant FN form where we mutated them to alanines, rendering them incompetent for oxidation (FN^ΔOxi^). As a control, we mutated the α5β1 and αv integrin binding site RGD to RGE (FN^ΔRGD^) thus blocking their binding through this site^44^. Western blot analysis from cells overexpressing these two constructs demonstrated that they are expressed and secreted to similar levels as that of cells over-expressing the full-length wild-type FN protein (FN^WT^) (Figure 4C-D, Figure S4E).

Considering that LOX treatment enhances FN clustering (Figure 2), we took into account that under classical adhesion assays with excess FN adsorbed to the surface (typically 10-20 µg/ml), stochastic clustering readily occurs. Thus, to avoid the possible interference of pre-adsorbed FN, we performed early adhesion assays of the cells expressing the FN variants directly on glass coverslips with no FN coating. We cultured the cells for 1 hour with serum-free media (to avoid plasma FN present in the serum from affecting the experiment) and quantified their areas to test whether the non-oxidizable FN variant inhibits initiation of cell adhesion. We find that the over-expression of FN^ΔRGD^ led to a significant reduction in cell area and β1 integrin expression, an integrin expressed in later stages of the initiation of cell adhesion, demonstrating an inhibition of the FN-integrin interactions. Notably, the over-expression of FN^ΔOxi^ led to a similar reduction in both properties as that observed with the over-expression of FN^ΔRGD^, demonstrating the role that these oxidation sites play in the initiation of FN-dependent integrin activation (Figure 4E-I).

To explore whether these mutations affect also FN fibrillogenesis, we first tested whether cells fibrilize their own secreted FN. To that end, we coated coverslips with Alexa 647-labelled FN and cultured MEF FN-GFP (in which GFP is knocked-in to both alleles of the FN gene^25^) for 48 hours, followed by imaging of the FN fibrils. All FN fibrils we observed were GFP labelled, demonstrating that cells use their own secreted FN in forming the initial fibrils (Figure S4F-G). Next, we cultured the cells expressing the FN variants on glass without any external FN coating for 24 hours in serum-free media, immunostained them for FN, and monitored FN fibrils. Cells over-expressing the FN^ΔRGD^ demonstrated a significant 27% reduction in the ratio between fibrillar FN vs. total FN; thus, even though integrin binding was perturbed in the lack of the RGD site, alternative interactions with αvβ3 integrin receptors could promote FN fibrillogenesis^45^, but to a lesser extent. Strikingly, cells over-expressing FN^ΔOxi^ displayed significantly less fibrillar FN (>62% reduction), despite containing the wild-type RGD site, demonstrating that the oxidation is a prerequisite for adhesion initiation (Figure 4J-M). As these cells still express endogenous wild-type FN, these results strongly demonstrate the critical and essential role played by members of the LOX-family enzymes in regulating FN fibrillogenesis, establishing that initiation of the process is an enzyme-rather than force-dependent process, in line with their ability to generate force-bearing stress fibers (Figure 4N). Similarly, when we used human dermal fibroblasts or HASMC infected with *shControl* lentiviral particles we found that they nicely formed FN fibrils when cultured directly on glass, but *shLOX* dermal fibroblasts and HASMC (Figure S5A) generated significantly less FN fibrils, even though they were cultured on a very stiff matrix and formed stress fibers (Figure 4O-S, S5B-E). To verify the specificity of the *shLOX* phenotypes, we knocked down LOX using two additional independent *shLOX* lentiviral constructs. We find that they result in similar phenotypes demonstrating that the phenotypes are specific to LOX (Figure S5F-I).

Our results thus far suggest that FN oxidation by lysyl oxidases is a prerequisite for FN fibrillogenesis, advocating that fibrillogenesis initiation is an enzyme-dependent rather than force dependent process. Should this be the case, then blocking lysyl oxidases’ enzymatic activities and reducing cellular forces via blebbistatin, a myosin II inhibitor, should lead to additive effects. Since the use of βAPN during adhesion initiation abrogates adhesion, we applied the inhibitor 30 minutes following cell seeding. We find that this treatment significantly inhibits long-term fibrillogenesis as measured 24 hours later. An additive effect was observed following the co-culturing of the cells with βAPN and blebbistatin (Figure S5J-N). Altogether, these findings further demonstrate the necessity for lysyl oxidases’ enzymatic activity in regulating FN fibrillogenesis, integrin activation and cell adhesion.

### FN oxidation promotes cancer cells colony formation in soft agar and migration

ECM organization and deposition play key roles in tumorigenesis. Since FN fibrillogenesis underlies these ECM properties and breast cancer progression is highly dependent on FN, we set to test the dependence of breast cancer cell migration and their ability to form colonies on FN oxidation. Towards that end, we stably expressed 67NR breast cancer cells with FN^WT^, FN^ΔRGD^ or FN^ΔOxi^ (Figure S4) and monitored the above properties. The soft agar colony formation assay serves as one of the hallmarks of carcinogenesis^46^ and has been shown to be dependent on FN^47^. Hence, we monitored the ability of these breast cancer cells to form colonies. We find that the colonies over-expressing either FN^ΔRGD^ or FN^ΔOxi^ were significantly smaller than those over-expressing FN^WT^ (Figure 5A-D).

**Figure 5.**
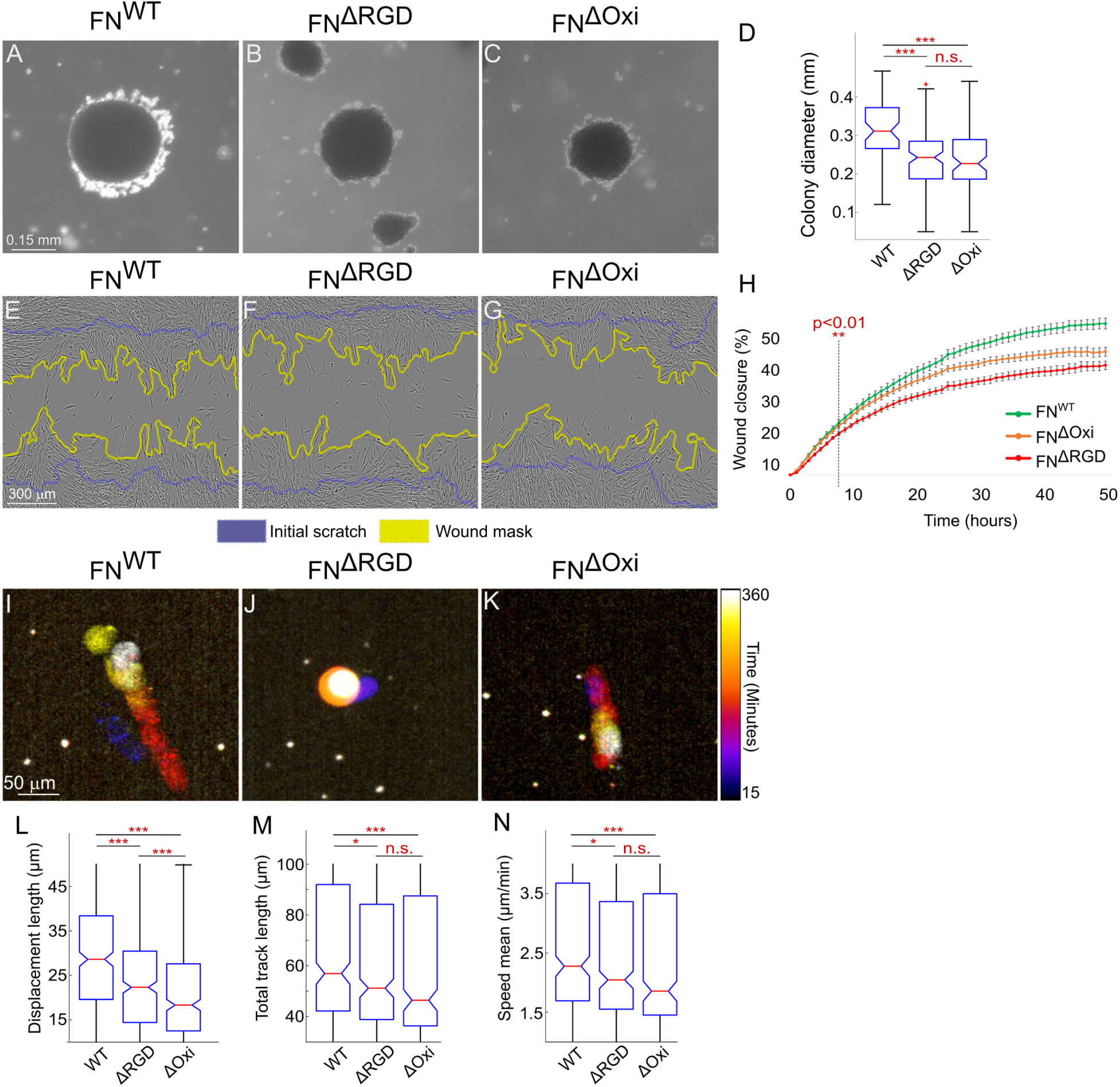
Mutations in LOX-specific oxidation sites within FN significantly impairs cancer cell characteristics. Representative images of the FN^WT^, FN^ΔRGD^ and FN^ΔOxi^ colonies formed in soft agar **(A-C)**. Quantification of colonies’ diameter show a significant decrease in FN^ΔOxi^ cells **(D, n>55)**. Representative images of the FN^WT^, FN^ΔRGD^ and FN^ΔOxi^ 67NR cell monolayers during wound healing assays **(E-G)**. Cumulative XY graph shows faster wound closure by FN^WT^ cells compared to FN^ΔOxi^ and FN^ΔRGD^ cells, demonstrating the significance of LOX-specific oxidation on cancer cell migration **(H, n>20)**. Representative time color-coded images generated from time-lapse videos tracking fluorescence-labeled nuclei FN^WT^, FN^ΔRGD^ and FN^ΔOxi^ 67NR cells **(I-K)**. Quantifications of single cell migration parameters show significant decrease in the measured parameters of cells infected with the FN mutants **(L-N, n>330)**. ***p<0.0001, **p<0.001, *p<0.01, n.s. = non-significant.

We next set to monitor the effects of these FN-derivatives on cell migration. Quantification of cell migration in a scratch assay over 48 hours (Figure 5E-H) as well as in a single cell migration assay over 6 hours (Figure 5I-N) demonstrated that, as above, the cells over-expressing either of the two FN mutant forms exhibited a reduced cell migration capacity.

Altogether, these results strongly demonstrate that the oxidation of FN on the 3 three lysine residues is essential for the cells ability to induce FN fibrillogenesis and as a result to modulate cell behavior

## Discussion

Our results reveal a - regulatory step essential for the formation of FN fibrils mediated by the various members of the lysyl oxidase enzyme family. We demonstrate that following lysyl oxidases inhibition (*shLOX* or βAPN) or mutation in the lysyl oxidases-specific oxidation sites within FN (mutant FN derivatives), FN fibrillogenesis is significantly perturbed, even in the background of endogenous wild-type FN. Conversely, we show that pretreatment of FN with recombinant lysyl oxidases promotes FN fibrillogenesis, cell adhesion and force generation in a FN-dependent manner, even on soft matrices.

We find that the enzymatic activity of lysyl oxidases leads to oxidation of specific lysine residues within FN, resulting in a structural change in the FN dimers thereby promoting early adhesion. We propose that through this structural change, a local increase in RGD density occurs, thus serving as a nucleator of FN fibrillogenesis. Following this nucleation step, integrin-mediated cell adhesion is favored, leading to increased force transmission to the matrix and the subsequent formation of bona-fide FN fibrils (Figure 6). This process could particularly be important for cells to overcome the soft embryonic environment through local secretion of LOX enzymes along migration routes, e.g., in the case of neural crest cells. Thus, similar to nucleation of cytoskeletal fibers’ polymerization (e.g., F-actin), which is regulated locally through the activity of nucleators (e.g., formins), initiation of FN fibrillogenesis also requires nucleation. As this nucleation step is an enzyme-dependent process, rather than a force-dependent one, these observations could bridge the observed discrepancies between the *in vivo* observations of FN fibrils in early embryos and in soft tissues, and the cell culture-based models which suggest that fibrillogenesis occurs primarily on stiff matrices.

**Figure 6.**
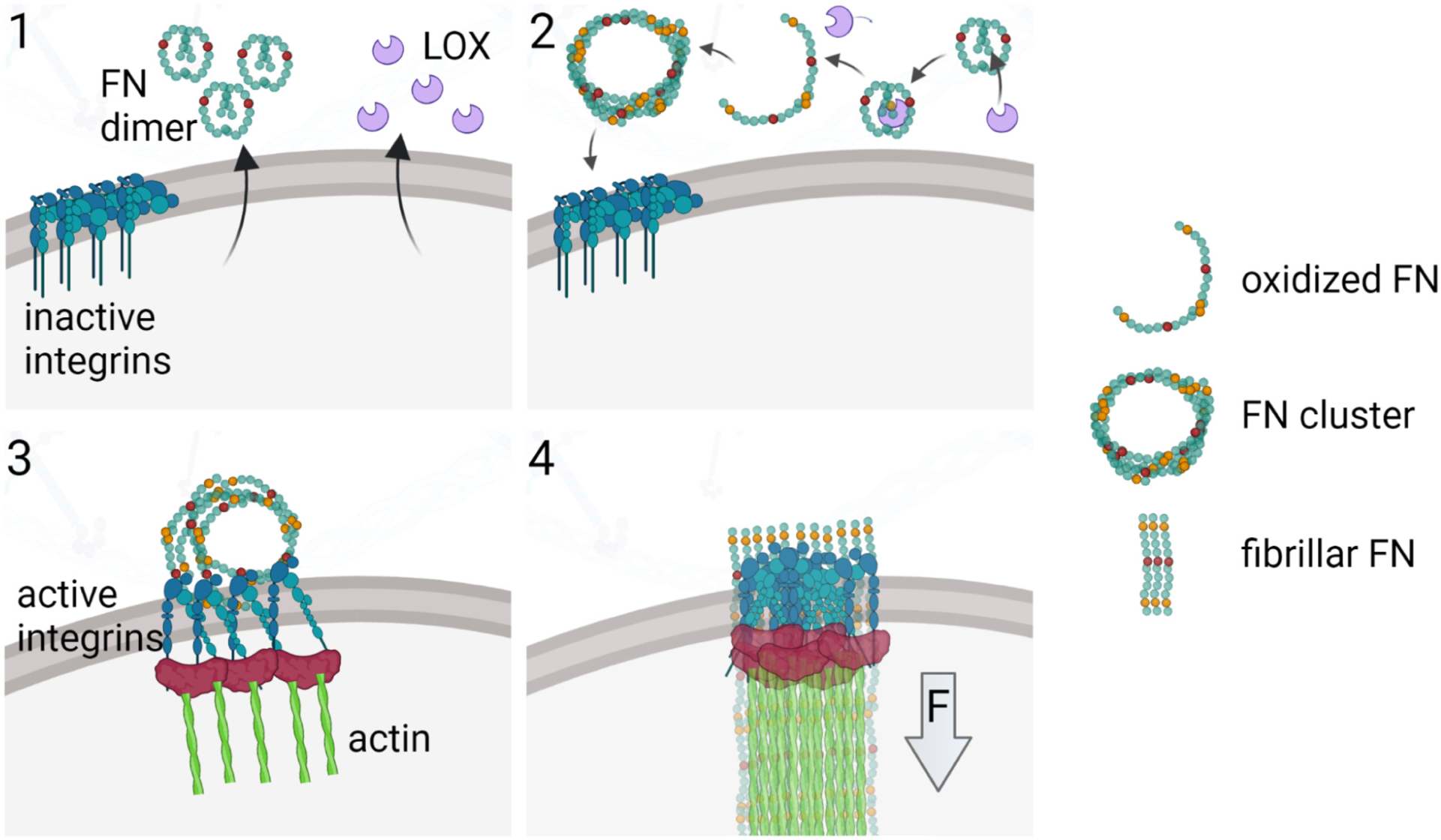
Model depicting the stages **(1-3)** required for generating FN clusters that act as nucleators for the force-induced fibrillogenesis step **(4)**. Created with BioRender.com.

Our results suggest that in the lack of LOX activity, cells perceive a soft environment even on rigid matrices. The association between LOX activity and rigidity perception highlight an alternative explanation to the observed association of lysyl oxidase upregulation and metastases^48^. Thus, cells initiate migration not only in response to general tissue stiffness but rather due to their local immediate environment, or the perception of their environment, even on soft matrices. As the majority of adhesion assays are carried out on FN coated matrices, we suggest that such conditions mask the LOX-dependent nucleation step and alter the perception of the immediate outside environment. Our results therefore imply that to identify and study the early steps of cell adhesion, cells must be cultured on matrices with either no FN or limiting FN levels. The ubiquitous expression of FN, and the ability of the various widely expressed members of the LOX family to oxidize it, suggest that the process that we have identified is general and is not specific to a distinct tissue or cell type. This process is thus expected to have important implications for our understanding of various matrix-dependent cellular functions in different tissues and contexts.

### Limitations of the study

Lysine oxidation is a key modification essential for the crosslinking of multiple ECM proteins, primarily collagens and elastin. Are FN fibrils also cross-linked in a similar manner or are the oxidations essential only for the nucleation and initiation of FN fibrillogenesis? Here, we focused mainly on in vitro studies during the initial steps of FN fibrillogenesis when the fibrils still hardly interact with other ECM components. Thus, we were not able to differentiate between the two options as this requires additional proteomic studies to be carried out also in vivo. Such in vivo studies, performed on distinct stages of embryonic development or various adult tissues, will further allow distinguishing whether FN oxidation is essential primarily on soft matrices as a cellular mechanism to overcome the lack of sufficient rigidity that initiates integrin activation or whether it is an essential step independent of matrix stiffness, questions that were not answered in this study.

## Acknowledgments

We would like to thank Mike Sheetz, Tom Schultheiss and Ella Preger-Ben Noon for critical reading and helpful discussions. H.W. is an incumbent of the David and Inez Myers Career Advancement Chair in Life Sciences. This research was supported by Israel Science Foundation grant 1738/17 (H.W.), Israel Science Foundation grant 111/18 (P.H.), Rappaport Family Foundation (H.W. and P.H.), National Heart, Lung, and Blood Institute grants R01 HL103920, R01 HL134935, R01 HL158049 (S.A.)

## Author contributions

S.M. designed the experiments and methodology, performed the experiments, analyzed the data, and wrote the manuscript. S.Z.E., E.N.B, R.A., H.Z., W.Y.B & R.K.A performed the experiments. D.S.D., O.L. & S.A. supervised the performed experiments and assisted in conceptualization of the study. H.W. & P.H. were responsible for conceptualization, supervision, funding acquisition, reviewing, writing and editing the manuscript.

## Inclusion and diversity

We support inclusive, diverse, and equitable conduct of research.

## STAR★Methods

### Resource Availability

#### Lead Contact

Further information and requests for resources and reagents should be directed to and will be fulfilled by the lead contact, Haguy Wolfenson.

#### Materials Availability

Following the completion of a material transfer agreement with the lead contact, all reagents generated in this study are available.

#### Data and Code Availability

- Mass-spectometry data have been deposited at PRIDE and are publicly available as of the date of publication. Accession numbers are listed in the key resources table. Microscopy data reported in this paper will be shared by the lead contact upon request.
- This paper does not report original code
- Any additional information required to reanalyze the data reported in this paper is available from the lead contact upon request.

#### Experimental Model and Study Participant Details

##### Mice maintenance and genotyping

All experiments involving mice were performed according to the relevant regulatory standards (Technion IACUC and national animal welfare laws, guidelines, and policies). All mice are housed in IVC’s (Techniplast) according to space requirements defined by the NRC. All rooms are set to have 22 ± 2 °C and humidity of 30–70%. HVAC parameters and light cycle are set by the computerized central system to full light of 10 hours, half-light of 2 hours and complete darkness of 12 hours. All mice were bred on a C57Bl/6 background purchased from Envigo (https://www.envigo.com). Embryonic day was staged according to Kaufmann, whereby the morning of the day in which a vaginal plug was observed was marked as E0.5. Mice embryos were secrefices at the embryonic stages of either E8, E9.5 AND E18.5. Mice were genotyped using PCR and the following primers: LoxL3: 5’- GCCAGGGTGAAGTGAAAGAC-3’; 3’-GATCTGGGATGCTGAAGACC-5’; 3’-GAACTTCGGAATAGGAACTTCG-5’. 300bp and 100bp represent wild-type and mutant PCR products, respectively.

##### Cell culture

MEF FN-GFP were described previously and were kindly provided by S. Astrof^49^. MEF Paxillin-GFP cells were kindly provided by B. Geiger (Weizmann institute of Science, Israel). Human Dermal fibroblasts (originated from male) were kindly provided by R. Shalom-Feuerstein (Technion, Israel). The three cell lines were cultured in DMEM (Sigma) with 10% FBS, 1% glutamine and 1% pen-strep (Biological Industries), or Optimem (Gibco, 31985070) with 1% glutamine and 1% pen-strep during experiments that require the absence of FN in the medium. HASMC (Sigma, 354-05A) were cultured with Smooth Muscle Cell Growth Medium 2 (PromoCell, C-22062), or Optimem (Gibco, 31985070) with 1% glutamine and 1% pen-strep during experiments that require the absence of FN in the medium.

#### Method Details

##### Embryos’ dissection, imaging and analysis

E9.5 and E18.5 embryos were dissected and underwent over-night fixation after somites were counted. Afterwards, whole mount staining was performed for the E9.5 embryos using FN and AP2α antibodies. Neural crest nuclei in the second branchial arch were then counted from confocal images. For analyses of FN organization in the somite borders, z-stack images were taken. We then used the trainable Weka segmentation plugin^50^ of ImageJ (NIH) software to identify FN fibrils in each stack. The area of the quantified fibrils was divided by the total somite area in each stack and the ratios were then averaged. For detection of cleft palate, E18.5 embryos underwent alcian blue and alizarin red staining.

##### Wholemount embryo staining

Embryos were incubated in 600 μl of blocking buffer (PBS, 0.01% Triton X-100 and 10% non-immune donkey serum (Sigma cat # D9663-10ml) overnight at 4 °C, then with 600 μl of blocking buffer containing primary antibodies for 4 days at 4 °C, with gentle rocking. The following primary antibodies were used: anti-Fn1 (Abcam, ab199056, 1:500), anti-AP2α (Developmental Studies Hybridoma Bank, DSHB-3B5, 1:200). After the incubation with primary antibodies, embryos were rinsed and washed with PBST (PBS with 0.05% Triton X-100) for 2 days at 4 °C, with gentle rocking.

Embryos were then incubated with 600 μl of blocking buffer containing DAPI (ThermoFisher, cat #D3571, 5 mg/ml stock diluted 1:1000) and secondary antibodies diluted 1:300 for 4 days at 4 °C. Alexa-labeled secondary antibodies were purchased from Invitrogen (donkey anti-mouse Alexa-647 cat # A31571 and donkey anti-rabbit Alexa-488 cat #A21206). After staining with DAPI and secondary antibodies, embryos were washed with PBST. Prior to imaging, embryos were dehydrated gradually with 25%, 50%, 75% of methanol diluted in dH2O for 10 minutes for each step, and then incubated with 100% methanol twice for 10 min each, and cleared in 50% BABB (BABB and 100% methanol at 1:1 (v/v)), and 100% BABB. BABB was generated by mixing benzyl alcohol (Sigma, #B1042) and benzyl benzoate (Sigma, #B6630) at 1:2 (v/v). Cleared embryos were placed between two coverslips (VWR #16004-312) separated by FastWells spacers (Grace Bio-Labs, #664113). Images were acquired using Nikon A1-HD25 inverted microscope equipped with a 20x water immersion objective (#MRD77200), numerical aperture 0.95, working distance 0.95 mm, and the NIS-Elements AR 5.11.01 64-bit software. Optical sections were collected at 0.55 µm intervals.

##### Mass spectrometry proteolysis

The proteins in the samples were suspended in 8M Urea, 100mM Ammonium bicarbonate and were reduced with 3mM DTT (60°C for 30 min), modified with 10mM iodoacetamide in 100mM ammonium bicarbonate (in the dark, room temperature for 30 min) and digested in 2M Urea, 25mM ammonium bicarbonate with modified trypsin (Promega) at a 1:50 enzyme-to-substrate ratio, overnight at 37oC.

##### Mass spectrometry analysis

The eluted peptides were desalted using C18 tips (Top-tip, Glygen) dried and re-suspended in 0.1% Formic acid. The peptides were resolved by reverse-phase chromatography on 0.075 × 180-mm fused silica capillaries (J&W) packed with Reprosil reversed phase material (Dr Maisch GmbH, Germany). The peptides were eluted with linear 30 minutes gradient of 5% to 28% acetonitrile with 0.1% formic acid in water, 15 minutes gradient of 28% to 95% acetonitrile with 0.1% formic acid in water and 15 minutes at 95% acetonitrile with 0.1% formic acid in water at flow rates of 0.15 μl/min. Mass spectrometry was performed by Q-Exactive plus mass spectrometer (Thermo) in a positive mode using repetitively full MS scan followed by collision induces dissociation (HCD) of the 10 most dominant ions selected from the first MS scan. The mass spectrometry data was analyzed using the MaxQuant software 1.5.2.8^51^ for peak picking and identification using the Andromeda search engine, searching against the human proteome from the Uniprot database with mass tolerance of 6 ppm for the precursor masses and 20 ppm for the fragment ions. Oxidation on Lys and Met and protein N-terminus acetylation were accepted as variable modifications and carbamidomethyl on cysteine was accepted as static modifications. Minimal peptide length was set to six amino acids and a maximum of two miscleavages was allowed. The data was quantified by label free analysis using the same software. Peptide- and protein-level false discovery rates (FDRs) were filtered to 1% using the target-decoy strategy.

For proteomic analysis of the ECM secreted by HASMC, shotgun proteomics identified the oxidized form on lysine 116, however at low abundance, which is not surprising given the noisy sample of various ECM proteins secreted by the cells. Therefore, we used targeted MS analysis using the known masses and retention times from the shotgun analyses that were done with LOXL2 and LOXL3. Using this targeted search, we identified the oxidation on site 116 (see Table S3).

##### Generation of mutant FN constructs

Point mutations were generated on a human fibronectin pMAX vector plasmid (Addgene, #120402) using the Q5 site-directed mutagenesis kit (BioLabs, #E05545). The mutagenesis procedure was performed using the following primers: Oxidation site 1: 5’-TGAGCGTCCTGCAGACTCCATG-3’, 3’- TAAGTGTCACCCACTCGG-5’; RGD: 5’- GCCGTGGAGAAAGCCCCGCAA-3’, 3’-CAGTGACAGCATACACAGTGATGGTATAATCAAC-5’; Oxidation site 2: 5’- TGATGATGGGGCGACATACCACG-3’, 3’-TAACACGTTGCCTCATGAG-5’; Oxidation site 3: 5’- ACAGTGGCAGGCGGAATATCTCG-3’, 3’- TCTCCTACGTGGTATGTC-5’. Next, mutated FN constructs were PCR amplified and ligated into BamH1+Xho1 cut NSPI vector using NEBuilder® HiFi DNA Assembly kit (BioLabs, #E5520).

##### Lenti viral infection

Plasmids were transfected into HEK 293-FT cells using CalFectin™ Mammalian Cell Transfection Reagent (SignaGen, SL100478). After 48 hours, conditioned medium was collected and used to infect MEF or HASMC cells. Plasmids used: Fibronectin mutant constructs, shLOX (Sigma, #1 - TRCN0000045991, #2 - TRCN00000293898, #3 - TRCN00000286463), shControl (Sigma, SHC016).

##### Spreading and soft gel experiments

Ibidi 8-well plates (Ibidi, 190814/1) were coated with human plasma FN (hFN, Sigma-Aldrich, FC010-10MG, 10 µg/ml) or laminin (Corning, 354232, 10 µg/ml) for 1 hour at 37⁰C. Afterwards, the samples were treated with human recombinant LOXL3 (RhLOXL3, R&D systems, 6069-AO) or LOXL2 (RhLOXL2, R&D systems, 2639-AO) for 5 hours (or sodium borate buffer 50 mM as control), with or without 200 mM βAPN (Sigma, A3134-25G) and then washed. Then, cells were seeded on the FN matrix and fixed at varying time points after spreading, followed by immunostaining. For soft gel experiments, Ibidi 8-well plates were covered with silicone gels (DOWSIL, CY 52-276 A&B), prepared at different rigidities that were previously calibrated and described^52^ and which we verified by rheometric measurements. For the majority of the experiments, FN was adsorbed to the elastomer via hydrophobic interactions^33^ by 1 hour incubation at 37⁰C. In the case of the cross-linking of FN to the gel, plasma treatment was done prior to FN coating following the incubation of the gel with (3-Aminopropyl) triethoxysilane (APTES, Sigma, 440140) for 5 minutes. Then, APTES was washed, and the gel was incubated with 25% glutaraldehyde for 1 hour. Eventually, the gel was coated with control or treated FN for 1 hour at 37⁰C and the same experiment and analysis was done. Images of the cells on the gels were acquired using an LSM800 confocal microscope (Zeiss) with Airyscan function and 20X or 63X objectives.

##### Adhesion, cell area and fibronectin analysis

Images were blindly and randomly chosen for acquisition. All of the adhesion and fibronectin fibrils analyses shown in this paper were performed using the trainable Weka segmentation plugin^53^ of ImageJ (NIH) software.

##### Spatial interaction analysis

Co-localization analysis between β1 integrin and LoxL3 was done using interaction analysis of Mosaic FIJI plugin for spatial interactions^26^. Briefly, mask of the β1 integrin nascent adhesions was extracted to mark the areas of potential interactions. Then, the average size of nascent adhesions was defined to identify them. Eventually, calculation of the co-localization was done by the plugin’s algorithm and the p-value of interaction was plotted for each image. The table shown in Figure 1 summarizes the p value distribution of the images that were taken either from the cell edge of from the cell center as comparison.

##### Immunofluorescence

Cells were fixed with 4% paraformaldehyde solution (36% stock, Sigma-Aldrich, 50-00-0, diluted with PBS) and permeabilized with 0.1% Triton X-100 in PBS (PBT 0.1%). Primary antibodies were incubated with the fixed cells over night at 4⁰C. After, secondary antibodies and phalloidin were incubated for 1 hour at room temperature. For dSTORM imaging, a second 10-minute fixation was performed at the end of the immunostaining protocol.

Antibodies that were used: anti-total Paxillin (Abcam, ab32084, 1:200); anti-FN (Abcam, ab2413, 1:500); anti-LoxL3 and anti-LoxL2 (produced and kindly gifted by G. Neufeld, Technion, 1:400); anti-β1 integrin (BD biosciences, 553715, 1:500). Phalloidin (Abcam, ab176757, 1:1000) and DAPI (Biolegend, 422801, 1:10,000) were also used.

##### Single cell migration assay

FN was treated with RhLOXL3, with or without βAPN as described above. Then, 24-well cell imaging plates (Eppendorf, EP-0030741005) were coated with the treated or untreated FN. Cells were incubated with 200 nM Sir-DNA (SpiroChrome, SC007) for 6 hours, and were then seeded. Two hours after seeding, a series of 64 images was taken (4 images/h) using the ImageXpress Micro Confocal High-Content Imaging system (Molecular Devices). Nuclei tracking analysis was performed using IMARIS software (Oxford instruments).

##### dSTORM imaging and analysis

Prior to imaging, FN was treated with RhLOXL3 as described above (molar ratio of 25:1) for 5 hours and was then diluted to 1 ng/ml. Then, #1.5 coverslips were coated with the diluted FN (treated, untreated or fresh FN without any incubation). The αFN (Abcam, ab2413) antibody was used to label FN, and the monoclonal antibody 9EG7 was used to label the extracellular domain of β1 integrin (BD biosciences, 553715, 1:500)^54^. AlexaFlour-647 and AlexaFlour-680 conjugated antibodies were then used as secondary antibodies, respectively. Fresh dSTORM buffer was prepared as previously described^55^. 2D dSTORM images were taken using a SAFe360 module (Abbelight Ltd, Cachan, France) coupled to Olympus IX83 inverted microscope using a 100x oil-immersion TIRF objective (NA 1.49) and 640 nm laser. The system is equipped with two sCMOS cameras PCO.panda4.2. A total of 15,000 frames at 50 ms exposure time were acquired and used for single-molecule detections to reconstruct a dSTORM image. Resulting coordinate tables and images were processed and analyzed using SAFe NEO software (Abbelight Ltd, Cachan, France).

3D dSTORM images were acquired in a similar manner with the following changes: LoxL3 was detected using anti-LoxL3 antibody (produced and kindly gifted by G. Neufeld, Technion, 1:100). Integrin β1 was detected using anti-activated CD29 (β1 integrin, BD biosciences, 553715, 1:100). For identification of the Z axis position of each single molecule an astigmatic lens was place in front of the cameras.

##### Pillar array preparation and imaging

Pillar fabrication was performed by pouring Polydimethylsiloxane (PDMS) (mixed at 10:1 with its curing agent, Sylgard 184; Dow Corning) into silicon molds (fabricated as previously described^36^) with holes at fixed depths and distances (center-to-center distance = 1 µm). The molds were then placed, face down, onto glass-bottom 35 mm dishes (#0 coverslip, Cellvis) which were incubated at 65°C for 12h to cure the PDMS. The molds were next peeled off while immersed in ethanol to prevent pillar collapse. On the day of the experiment, FN or laminin were treated with rhLOXL3 for 5 hours (or sodium borate buffer 50 mM as control). Then, the pillars were coated with the treated or untreated FN for 1 hour at 37°c. Next, the buffer was replaced to HBSS + HEPES 20 mM (Biological industries), pH 7.2.

Pillar bending stiffness (*k*) was calculated by Euler–Bernoulli beam theory as described:

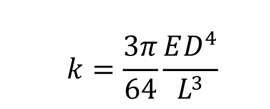

Where D and L are the diameter and length of the pillar, respectively, and E is the Young’s modulus of the PDMS (= 2 MPa).

Cells were resuspended with the HBSS/HEPES buffer and then spread on the FN-coated pillars. Time-lapse imaging of cells spreading on the pillars was performed using an inverted microscope (Leica DMIRE2) at 37°C using a 63x 1.4 NA oil immersion objective. Brightfield images were recorded every 2 seconds with a Retiga EXi Fast 1394 CCD camera (QImaging). The microscope and camera were controlled by Micromanager software^56^. For each cell, a movie of 10-20 minutes was recorded. To minimize photo-damage to the cells, a 600 nm longpass filter was inserted into the illumination path.

##### Pillar displacement analysis

Pillar tracking was performed using the Nanotracking plugin of ImageJ, as described previously^34^. In short, the cross-correlation between the pillar image in every frame of the movie and an image of the same pillar from the first frame of the movie was calculated, and the relative x-and y-position of the pillar in every frame of the movie was obtained. To consider only movements of pillar from their zero-position, we only analyzed pillars that at the start of the movie were not in contact with the cell and that during the movie the cell edge reached to them. Drift correction was performed using data from pillars far from any cell in each movie. For each pillar, the displacement curve was generated by Matlab (MathWorks). Analyses of the catch-and-release events was performed by the ‘findpeaks’ function in Matlab, considering all the peaks above noise level with a minimum width of 3 frames.

##### Western Blot

Protein lysates were harvested from the cell lines using a lysis buffer (Tris 10 mM pH7, 2 nM EDTA, 1% NP-40, 0.1% DOC, 0.2 mM AEBSF). The lysates were loaded onto SDS-PAGE. Gels were transferred to nitrocellulose membrane that was blocked for 1 hour (5% BSA, 0.1% Tween in TBS). The membranes were then probed with the primary antibody over night at 4 ⁰C. Secondary antibody was added for 2 hours at room temperature. Antibodies used for this method: anti-Lox (58135, Cell Signaling); anti-P97 (kindly provided by A. Stanhill, Technion); anti-Fibronectin (Abcam, ab23750).

##### Neural crest explants

Neural tube explants were carried out essentially as previously described^57^. Briefly, hindbrain regions of E8-8.5 embryos were dissected and the neural primordia consisting of the pre-migratory neural crest cells was isolated from surrounding tissues using fine dissecting pins. The neural tubes were then cultured on FN (50 µg/ml) coated dishes and incubated for 16 hours at 37⁰C in DMEM supplemented with 10% fetal bovine serum, 1% 1M HEPES, 1% Pen Strep, and 1% L-glutamate (Gibco, USA). Explants were then fixed with 4% PFA and immunofluorescently stained.

##### Soft agar assay

Protocol was done as previously described^58^. Briefly, the lower layer of 0.5% agarose was prepared using pre-melted 1% agarose and 2x concentrated culture medium and distributed in a 6-well dish. The dish was then incubated for 30 minutes at 4⁰C to coagulate the agar. After, the upper layer of 0.3% agarose was prepared using pre-melted 0.6% agarose and 2x concentrated culture medium. 67NR cells were suspended in the upper layer (4000 cells/well) and were distributed above the 0.5% agarose layer that was previously prepared. The 6-well plate was incubated again to coagulate the agar. Then, 300ul of culture medium was added on top of the agar every 3 days. Cells were fixed after 17 days using 10% ethanol in PBS and stained with crystal violet. 4x images of the colonies were taken using light microscope and colonies’ diameter was measured.

##### Wound healing assay

Scratch assay were performed, and cell migration was monitored by the automated Incucyte ZOOM (Sartorius, Gottingen, Germany). 67NR cells infected with the FN mutant plasmids were seeded in an Essen Imagelock 96-well plate at the density of 1×60^4^ cells/well and were maintained over night to reach 100% confluency. A wound was made in the cell monolayer using a 96-well wound-maker tool with PTFE pin tips (Sartorius) according to the manufacturer’s instructions. Afterwards, the plate was inserted into the Incucyte ZOOM and incubated in 5% CO2 at 37°C for 53 hours. Images were captured at 1-hour intervals, and data were analyzed using the IncuCyte Zoom software (Sartorius). Relative wound density (RWD), which measures the relative cell density in the wounded and non-wounded areas at each time point, was used to assess the rate of cell migration.

#### Quantification and Statistical Analysis

Matlab was used for statistical analysis and graph plotting. The data’s normality and homogeneity of variances were checked for each experiment and then a parametric or non-parametric statistical test was performed for each data set. In the case of two groups that are normally distributed and have similar variances, T-test was performed. Otherwise, the non-parametric Mann Whitney U test was performed. In case of three or four groups that are normally distributed, have similar variances and have no multicollinearity, ANOVA test was performed. Otherwise, non-parametric Kruskal-Wallis test was performed. In the case of histograms, two-sample Kolmogorov-Smirnov (KS) test was performed. For all of the boxplots shown in the paper, the red central mark represents the median, the bottom and the top edges of the box represents the 25th and 75th percentiles, respectively, and the black top and bottom represent the minimal and maximal values that are not outliers, respectively. Outliers are represented as red crosses. P values represented as ***p<0.0001, **p<0.001, *p<0.05, n.s. = non-significant.

#### Key resources table

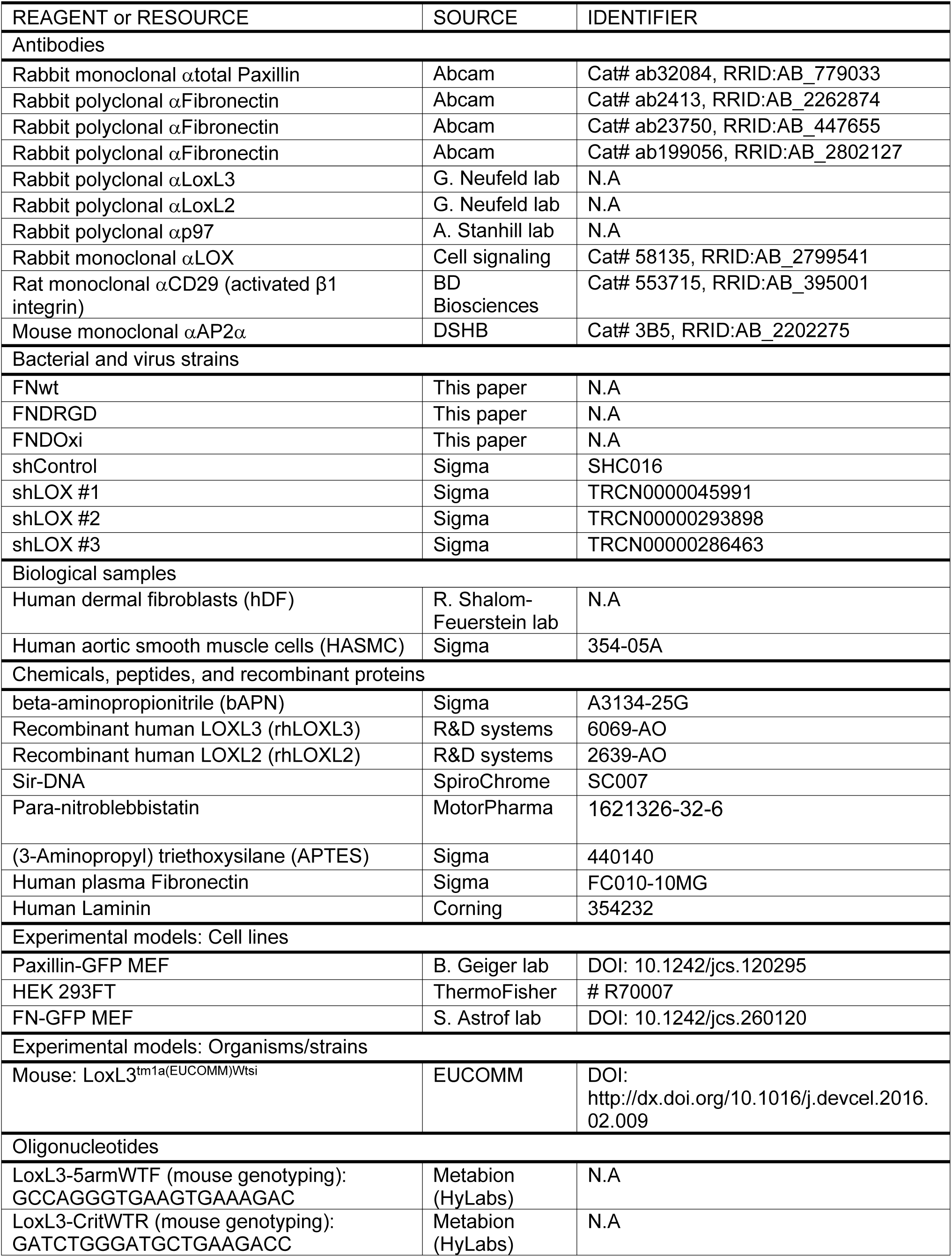
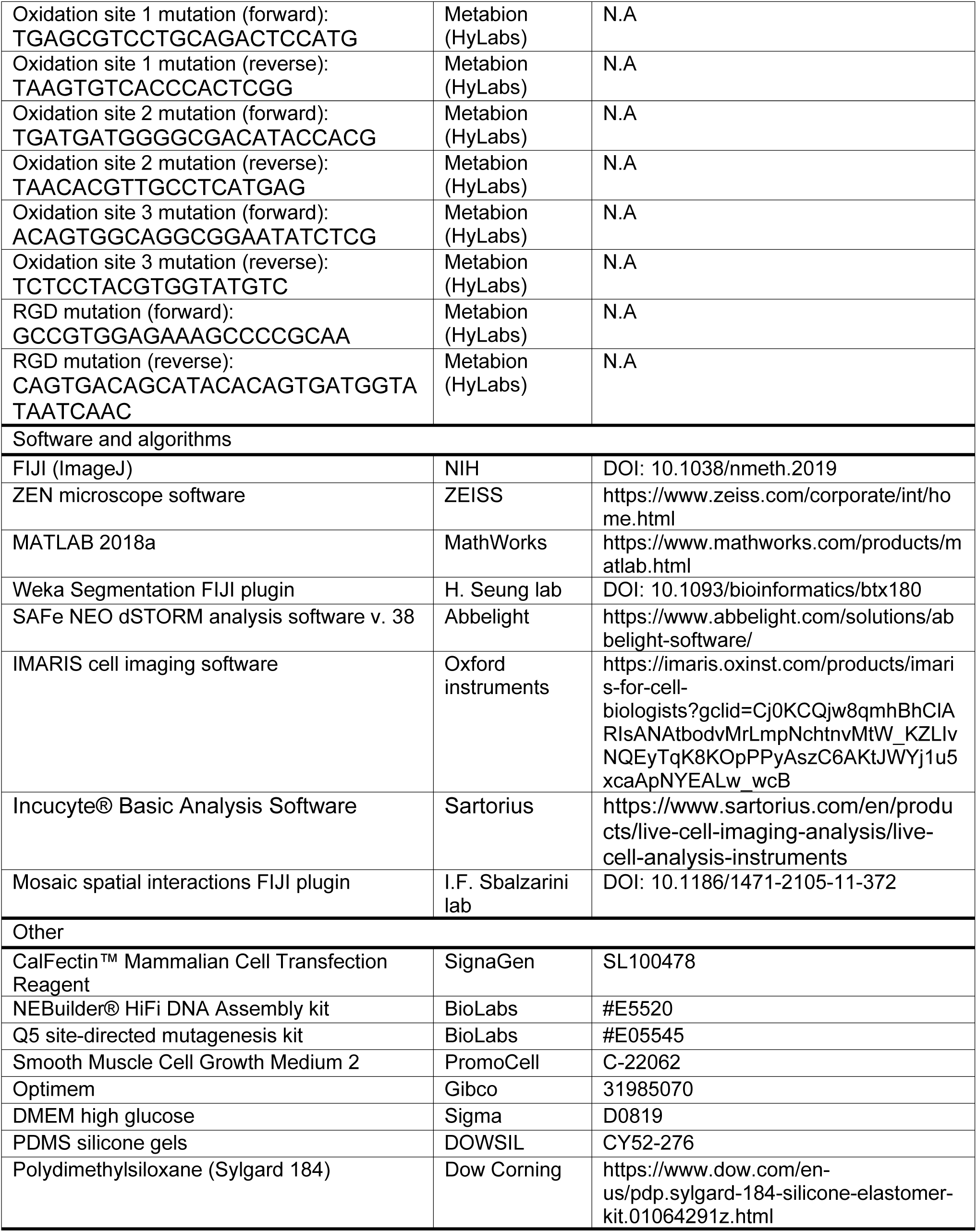

## Supplemental tables titles

**Table S1:** LOXL3-FN1 specific peptides

**Table S2:** LOXL2-FN1 specific peptides

**Table S3:** HASMC identified oxidations

## References

1. Bonnans, C., Chou, J., and Werb, Z. (2014). Remodelling the extracellular matrix in development and disease. Nature Reviews in Molecular Cell Biology 15, 786–801. 10.1038/nrm3904.

2. Humphrey, J.D., Dufresne, E.R., and Schwartz, M.A. (2014). Mechanotransduction and extracellular matrix homeostasis. Nature Reviews in Molecular Cell Biology 15, 802–812. 10.1038/nrm3896.

3. Schwarzbauer, J.E., and DeSimone, D.W. (2011). Fibronectins, their fibrillogenesis, and in vivo functions. Cold Spring Harb Perspect Biol 3, 1–19. 10.1101/cshperspect.a005041.

4. Pankov, R., and Yamada, K.M. (2002). Fibronectin at a glance. J Cell Sci 115, 3861–3863. 10.1242/jcs.00059.

5. Gold, L.I., and Pearlstein, E. (1980). Fibronectin–collagen binding and requirement during cellular adhesion. Biochemical Journal 186, 551–559. 10.1042/BJ1860551.

6. Sabatier, L., Chen, D., Fagotto-Kaufmann, C., Hubmacher, D., McKee, M.D., Annis, D.S., Osher, D.F., and Reinhardt, D.P. (2009). Fibrillin assembly requires fibronectin. Mol Biol Cell 20, 846–858. 10.1091/mbc.e08-08-0830.

7. Pereira, M., Rybarczyk, B.J., Odrljin, T.M., Hocking, D.C., Sottile, J., and Simpson-Haidaris, P.J. (2002). The incorporation of fibrinogen into extracellular matrix is dependent on active assembly of a fibronectin matrix. J Cell Sci 115, 609–617. 10.1242/JCS.115.3.609.

8. Chung, C.Y., and Erickson, H.P. (1997). Glycosaminoglycans modulate fibronectin matrix assembly and are essential for matrix incorporation of tenascin-C. J Cell Sci 110, 1413–1419. 10.1242/JCS.110.12.1413.

9. Singh, P., Carraher, C., and Schwarzbauer, J.E. (2010). Assembly of Fibronectin Extracellular Matrix. http://dx.doi.org/10.1146/annurev-cellbio-100109-10402026, 397–419. 10.1146/ANNUREV-CELLBIO-100109-104020.

10. Früh, S.M., Schoen, I., Ries, J., and Vogel, V. (2015). Molecular architecture of native fibronectin fibrils. Nat Commun 6. 10.1038/ncomms8275.

11. Zhong, C., Chrzanowska-Wodnicka, M., Brown, J., Shaub, A., Belkin, A.M., and Burridge, K. (1998). Rho-mediated contractility exposes a cryptic site in fibronectin and induces fibronectin matrix assembly. J Cell Biol 141, 539–551. 10.1083/jcb.141.2.539.

12. Antia, M., Baneyx, G., Kubow, K.E., and Vogel, V. (2008). Fibronectin in aging extracellular matrix fibrils is progressively unfolded by cells and elicits an enhanced rigidity response. Faraday Discuss 139, 229–249. 10.1039/b718714a.

13. Cao, L., Nicosia, J., Larouche, J., Zhang, Y., Bachman, H., Brown, A.C., Holmgren, L., and Barker, T.H. (2017). Detection of an Integrin-Binding Mechanoswitch within Fibronectin during Tissue Formation and Fibrosis. ACS Nano 11, 7110. 10.1021/ACSNANO.7B02755.

14. Carraher, C.L., and Schwarzbauer, J.E. (2013). Regulation of matrix assembly through rigidity-dependent fibronectin conformational changes. J Biol Chem 288, 14805–14814. 10.1074/jbc.M112.435271.

15. Baneyx, G., Baugh, L., and Vogel, V. (2002). Fibronectin extension and unfolding within cell matrix fibrils controlled by cytoskeletal tension. Proc Natl Acad Sci U S A 99, 5139–5143. 10.1073/PNAS.072650799.

16. Wolfenson, H., Lavelin, I., and Geiger, B. (2013). Dynamic Regulation of the Structure and Functions of Integrin Adhesions. Dev Cell 24, 447–458. 10.1016/J.DEVCEL.2013.02.012.

17. Zamir, E., Katz, M., Posen, Y., Erez, N., Yamada, K.M., Katz, B.-Z., Lin, S., Lin, D.C., Bershadsky, A., Kam, Z., et al. (2000). Dynamics and segregation of cell-matrix adhesions in cultured fibroblasts. Nat Cell Biol 2. 10.1038/35008607.

18. Carraher, C.L., and Schwarzbauer, J.E. (2013). Regulation of matrix assembly through rigidity-dependent fibronectin conformational changes. J Biol Chem 288, 14805–14814. 10.1074/jbc.M112.435271.

19. Dzamba, B.J., Jakab, K.R., Marsden, M., Schwartz, M.A., and DeSimone, D.W. (2009). Cadherin Adhesion, Tissue Tension, and Noncanonical Wnt Signaling Regulate Fibronectin Matrix Organization. Dev Cell 16, 421–432. 10.1016/J.DEVCEL.2009.01.008.

20. Agero, U., Glazier, J.A., and Hosek, M. (2010). Bulk elastic properties of chicken embryos during somitogenesis. Biomed Eng Online 9, 1–16. 10.1186/1475-925X-9-19.

21. Marrese, M., Antonovaite, N., Nelemans, B.K.A., Smit, T.H., and Iannuzzi, D. (2019). Micro-indentation and optical coherence tomography for the mechanical characterization of embryos: Experimental setup and measurements on chicken embryos. Acta Biomater 97, 524–534. 10.1016/J.ACTBIO.2019.07.056.

22. Rifes, P., and Thorsteinsdóttir, S. (2012). Extracellular matrix assembly and 3D organization during paraxial mesoderm development in the chick embryo. Dev Biol 368, 370–381. 10.1016/j.ydbio.2012.06.003.

23. de Almeida, P.G., Pinheiro, G.G., Nunes, A.M., Gonçalves, A.B., and Thorsteinsdóttir, S. (2016). Fibronectin assembly during early embryo development: A versatile communication system between cells and tissues. Developmental Dynamics 245, 520–535. 10.1002/dvdy.24391.

24. Kraft-Sheleg, O., Zaffryar-Eilot, S., Genin, O., Yaseen, W., Soueid-Baumgarten, S., Kessler, O., Smolkin, T., Akiri, G., Neufeld, G., Cinnamon, Y., et al. (2016). Localized LoxL3-Dependent Fibronectin Oxidation Regulates Myofiber Stretch and Integrin-Mediated Adhesion. Dev Cell 36, 550–561. 10.1016/j.devcel.2016.02.009.

25. Tomer, D., Arriagada, C., Munshi, S., Alexander, B.E., French, B., Vedula, P., Caorsi, V., House, A., Guvendiren, M., Kashina, A., et al. (2022). A new mechanism of fibronectin fibril assembly revealed by live imaging and super-resolution microscopy. J Cell Sci 135, jcs260120. 10.1242/jcs.260120.

26. Helmuth, J.A., Paul, G., and Sbalzarini, I.F. (2010). Beyond co-localization: inferring spatial interactions between sub-cellular structures from microscopy images. BMC Bioinformatics 11, 372. 10.1186/1471-2105-11-372.

27. Mittal, A., Pulina, M., Hou, S.Y., and Astrof, S. (2010). Fibronectin and integrin alpha 5 play essential roles in the development of the cardiac neural crest. Mech Dev 127, 472–484. 10.1016/J.MOD.2010.08.005.

28. Wang, X., and Astrof, S. (2016). Neural crest cell-autonomous roles of fibronectin in cardiovascular development. Development 143, 88–100. 10.1242/DEV.125286/256986.

29. Matsumoto, T., Sugita, S., and Nagayama, K. (2016). Tensile Properties of Smooth Muscle Cells, Elastin, and Collagen Fibers BT - Vascular Engineering: New Prospects of Vascular Medicine and Biology with a Multidiscipline Approach. In, K. Tanishita and K. Yamamoto, eds. (Springer Japan), pp. 127–140. 10.1007/978-4-431-54801-0_7.

30. S., L.V., M., H.C., P., H.E., Nikkola, C., Ignaty, L., G., L.C., J., B.A., Dana, V., null, null, P., M.R., et al. (2016). Loss of function mutation in LOX causes thoracic aortic aneurysm and dissection in humans. Proceedings of the National Academy of Sciences 113, 8759–8764. 10.1073/pnas.1601442113.

31. Sampath Narayanan, A., Siegel, R.C., and Martin, G.R. (1972). On the inhibition of lysyl oxidase by β-aminopropionitrile. Biochem Biophys Res Commun 46, 745–751. https://doi.org/10.1016/S0006-291X(72)80203-1.

32. Trappmann, B., Gautrot, J.E., Connelly, J.T., Strange, D.G., Li, Y., Oyen, M.L., Cohen Stuart, M.A., Boehm, H., Li, B., Vogel, V., et al. (2012). Extracellular-matrix tethering regulates stem-cell fate. Nat Mater 11, 742. 10.1038/nmat3339.

33. Missirlis, D., Haraszti, T., Heckmann, L., and Spatz, J.P. (2020). Substrate Resistance to Traction Forces Controls Fibroblast Polarization. Biophys J 119, 2558–2572. 10.1016/j.bpj.2020.10.043.

34. Wolfenson, H., Meacci, G., Liu, S., Stachowiak, M.R.M.R., Iskratsch, T., Ghassemi, S., Roca-Cusachs, P., O’Shaughnessy, B., Hone, J., and Sheetz, M.P.M.P. (2016). Tropomyosin controls sarcomere-like contractions for rigidity sensing and suppressing growth on soft matrices. Nat Cell Biol 18, 33–42. 10.1038/ncb3277.

35. Feld, L., Kellerman, L., Mukherjee, A., Livne, A., Bouchbinder, E., and Wolfenson, H. (2020). Cellular contractile forces are nonmechanosensitive. Sci Adv 6. 10.1126/SCIADV.AAZ6997.

36. Ghassemi, S., Meacci, G., Liu, S., Gondarenko, A.A., Mathur, A., Roca-Cusachs, P., Sheetz, M.P., and Hone, J. (2012). Cells test substrate rigidity by local contractions on submicrometer pillars. Proc Natl Acad Sci U S A 109, 5328–5333. 1119886109 [pii] 10.1073/pnas.1119886109.

37. Wolfenson, H., Meacci, G., Liu, S., Stachowiak, M.R.M.R., Iskratsch, T., Ghassemi, S., Roca-Cusachs, P., O’Shaughnessy, B., Hone, J., and Sheetz, M.P.M.P. (2016). Tropomyosin controls sarcomere-like contractions for rigidity sensing and suppressing growth on soft matrices. Nat Cell Biol 18, 33–42. 10.1038/ncb3277.

38. Ghibaudo, M., Saez, A., Trichet, L., Xayaphoummine, A., Browaeys, J., Silberzan, P., Buguin, A., and Ladoux, B. (2008). Traction forces and rigidity sensing regulate cell functions. Soft Matter 4, 1836. 10.1039/b804103b.

39. Jiang, G., Giannone, G., Critchley, D.R., Fukumoto, E., and Sheetz, M.P. (2003). Two-piconewton slip bond between fibronectin and the cytoskeleton depends on talin. Nature 424, 334–337. 10.1038/nature01805 nature01805 [pii].

40. Schvartzman, M., Palma, M., Sable, J., Abramson, J., Hu, X., Sheetz, M.P., and Wind, S.J. (2011). Nanolithographic control of the spatial organization of cellular adhesion receptors at the single-molecule level. Nano Lett 11, 1306–1312. 10.1021/nl104378f.

41. Cavalcanti-Adam, E.A., Volberg, T., Micoulet, A., Kessler, H., Geiger, B., and Spatz, J.P. (2007). Cell spreading and focal adhesion dynamics are regulated by spacing of integrin ligands. Biophys J 92, 2964– 2974. 10.1529/biophysj.106.089730.

42. Vallet, S.D., Berthollier, C., Salza, R., Muller, L., and Ricard-Blum, S. (2020). The Interactome of Cancer-Related Lysyl Oxidase and Lysyl Oxidase-Like Proteins. Cancers 2021, Vol. 13, Page 71 13, 71. 10.3390/CANCERS13010071.

43. Parsons, J.T., Horwitz, A.R., and Schwartz, M.A. (2010). Cell adhesion: integrating cytoskeletal dynamics and cellular tension. Nature Reviews in Molecular Cell Biology 11, 633–643. 10.1038/nrm2957.

44. Takahashi, S., Leiss, M., Moser, M., Ohashi, T., Kitao, T., Heckmann, D., Pfeifer, A., Kessler, H., Takagi, J., Erickson, H.P., et al. (2007). The RGD motif in fibronectin is essential for development but dispensable for fibril assembly. Journal of Cell Biology 178, 167–178. 10.1083/JCB.200703021.

45. Takahashi, S., Leiss, M., Moser, M., Ohashi, T., Kitao, T., Heckmann, D., Pfeifer, A., Kessler, H., Takagi, J., Erickson, H.P., et al. (2007). The RGD motif in fibronectin is essential for development but dispensable for fibril assembly. Journal of Cell Biology 178, 167–178. 10.1083/JCB.200703021.

46. Du, F., Zhao, X., and Fan, D. (2017). Soft Agar Colony Formation Assay as a Hallmark of Carcinogenesis. Bio Protoc 7, e2351. 10.21769/BioProtoc.2351.

47. Gupta, R.K., and Johansson, S. (2013). Fibronectin Assembly in the Crypts of Cytokinesis-Blocked Multilobular Cells Promotes Anchorage-Independent Growth. PLoS One 8, e72933. 10.1371/JOURNAL.PONE.0072933.

48. Amendola, P.G., Reuten, R., and Erler, J.T. (2019). Interplay Between LOX Enzymes and Integrins in the Tumor Microenvironment. Cancers (Basel) 11. 10.3390/CANCERS11050729.

49. Tomer, D., Arriagada, C., Munshi, S., Alexander, B.E., French, B., Vedula, P., Caorsi, V., House, A., Guvendiren, M., Kashina, A., et al. (2022). A new mechanism of fibronectin fibril assembly revealed by live imaging and super-resolution microscopy. bioRxiv, 2020.09.09.290130. 10.1101/2020.09.09.290130.

50. Chermnykh, E., Kalabusheva, E., and Vorotelyak, E. (2018). Extracellular matrix as a regulator of epidermal stem cell fate. Int J Mol Sci 19. 10.3390/ijms19041003.

51. Rgen Cox, J., Hein, M.Y., Luber, C.A., Paron, I., Nagaraj, N., and Mann, M. (2014). Accurate Proteome-wide Label-free Quantification by Delayed Normalization and Maximal Peptide Ratio Extraction, Termed MaxLFQ* □ S Technological Innovation and Resources. Molecular & Cellular Proteomics 13, 2513–2526. 10.1074/mcp.

52. Ou, G., Thakar, D., Tung, J.C., Miroshnikova, Y.A., Dufort, C.C., Gutierrez, E., Groisman, A., and Weaver, V.M. (2016). Visualizing mechanical modulation of nanoscale organization of cell-matrix adhesions. Integr Biol (Camb) 8, 795–804. 10.1039/c6ib00031b.

53. Chermnykh, E., Kalabusheva, E., and Vorotelyak, E. (2018). Extracellular matrix as a regulator of epidermal stem cell fate. Int J Mol Sci 19. 10.3390/ijms19041003.

54. Mandicourt, G., Iden, S., Ebnet, K., Aurrand-Lions, M., and Imhof, B.A. (2007). JAM-C regulates tight junctions and integrin-mediated cell adhesion and migration. Journal of Biological Chemistry 282, 1830– 1837. 10.1074/jbc.M605666200.

55. Nahidiazar, L., Agronskaia, A. V., Broertjes, J., Van Broek, B. Den, and Jalink, K. (2016). Optimizing imaging conditions for demanding multi-color super resolution localization microscopy. PLoS One 11, 1–18. 10.1371/journal.pone.0158884.

56. Edelstein, A., Amodaj, N., Hoover, K., Vale, R., and Stuurman, N. (2010). Computer control of microscopes using microManager. Curr Protoc Mol Biol 92, 14.20.1–14.20.17. 10.1002/0471142727.mb1420s92.

57. Kalev-Altman, R., Hanael, E., Zelinger, E., Blum, M., Monsonego-Ornan, E., and Sela-Donenfeld, D. (2020). Conserved role of matrix metalloproteases 2 and 9 in promoting the migration of neural crest cells in avian and mammalian embryos. The FASEB Journal 34, 5240–5261. https://doi.org/10.1096/fj.201901217RR.

58. Borowicz, S., Van Scoyk, M., Avasarala, S., Karuppusamy Rathinam, M.K., Tauler, J., Bikkavilli, R.K., and Winn, R.A. (2014). The soft agar colony formation assay. J Vis Exp, e51998. 10.3791/51998.

