## Supplementary Material for "Initiation of fibronectin fibrillogenesis is an enzyme-dependent process"

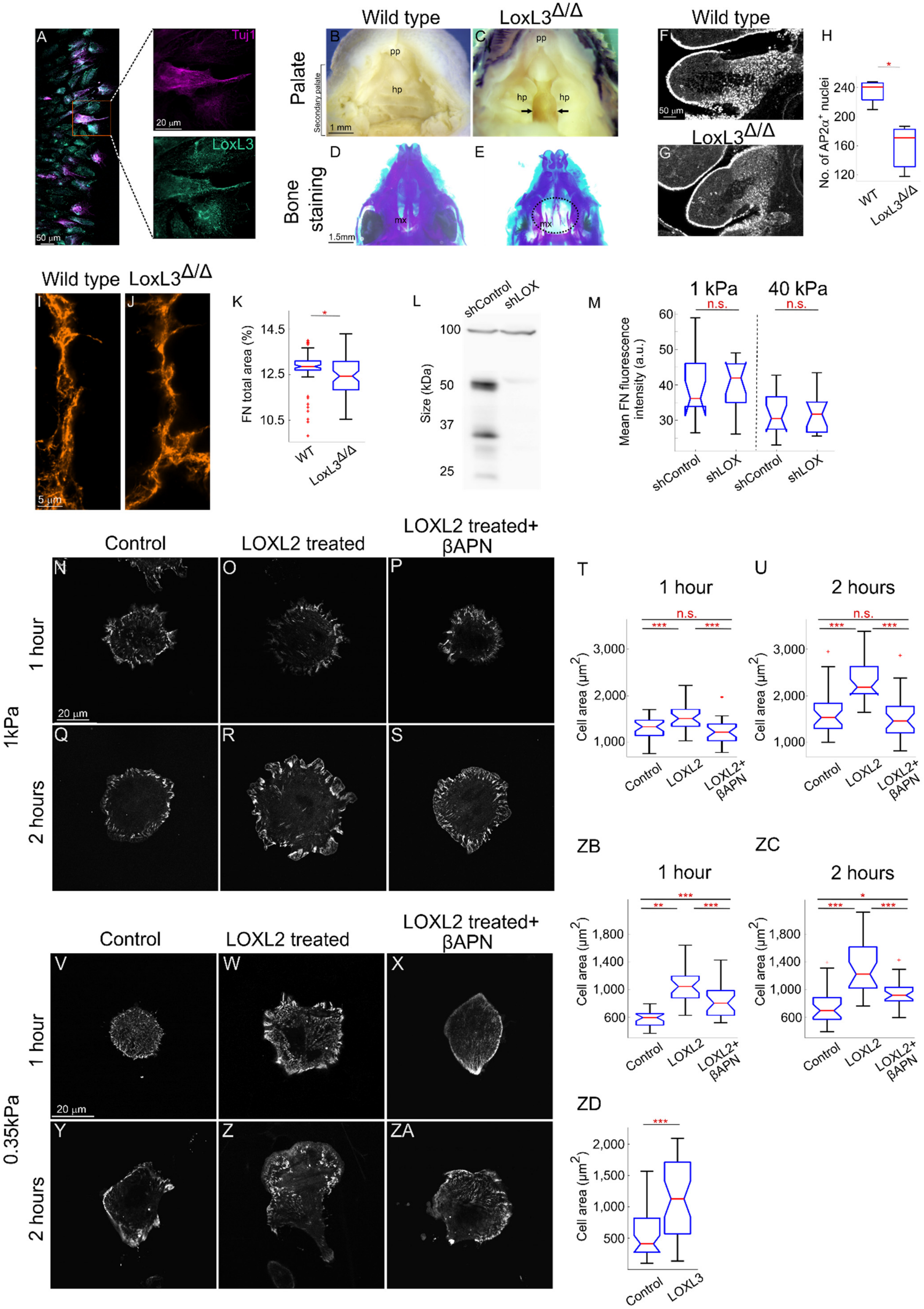

A
Tuj1
LoxL3
50 µm
20 µm
Wild type
LoxL3Δ/Δ
Palate
Secondary palate
B
C
pp
hp
1 mm
Bone staining
D
E
mx
1.5mm
F
G
50 µm
H
No. of AP2α+ nuclei
240
200
160
120
WT
I
J
5 µm
K
FN total area (%)
14.5
12.5
10.5
L
shControl
shLOX
Size (kDa)
100
50
37
25
M
1 kPa
40 kPa
n.s.
Mean FN fluorescence intensity (a.u.)
60
50
40
30
Control
LOXL2 treated
LOXL2 treated+ βAPN
N
O
P
Q
R
S
1 hour
2 hours
1kPa
T
U
Cell area (µm2)
3,000
2,000
1,000
LOXL2
LOXL2+ βAPN
V
W
X
Y
Z
ZA
0.35kPa
ZB
ZC
1,800
1,400
1,000
600
ZD
2,000
1,500
500
LOXL3

**Figure S1. Lysyl oxidases alterations affect cell behavior, Related to Figure 1.** Immunofluorescence staining of cranial neural tube explants obtained from E8.5 embryos to analyse expression of LoxL3 in Tuj1-expressing neural crest cells (A). Ventral view of the maxilla of newborn *wild-type* and *LoxL3*$^{\Delta/\Delta}$ mouse with cleft palate (B-C). The palate consists of the primary palate (pp) and the secondary palate, with the hard palate (hp). *LoxL3*$^{\Delta/\Delta}$ mouse showing complete cleft palate (arrows in C). Ventral view of the cranial base of E18.5 *wild-type* and *LoxL3*$^{\Delta/\Delta}$ mice (D-E) stained with alizarin red and alcian blue marking mineralized bone and cartilage, respectively. Maxilla (mx) is highlighted (dashed oval in E). Ap2α immunostaining of the second branchial arch artery of *wild-type* and *LoxL3*$^{\Delta/\Delta}$ 29-30 somites mouse embryos (F-G). Quantification of the number of nuclei shows a significant reduction of Ap2α positive nuclei in *LoxL3*$^{\Delta/\Delta}$ embryos (H, n=4 of each genotype). FN immunostaining of the border between the first and second somites of *wild-type* and *LoxL3*$^{\Delta/\Delta}$ 26 somites embryos (I-J). Quantification shows a significant reduction in the total FN area in *LoxL3*$^{\Delta/\Delta}$ somite border (K, n=2 of each genotype). Western blot for LOX from HASMC lysates (L). Quantification of the mean fluorescence intensity in samples immunostained for FN showing no significant difference in secreted FN levels between shControl and shLOX cells (M, n=15 images of each condition). Pax-GFP on sham- and LOXL2-treated FN with or without βAPN, seeded on 1kPa (N-S) or 0.25kPa (V-ZA) gels and fixed after 1 hour (N-P, V-X) and 2 hours (Q-S, Y-ZA). Quantification of cell areas show significant differences between cells seeded on treated FN regardless of rigidities (T-U, ZB-ZC, n>28). Quantification of cell areas of cells seeded on FN that is covalently cross-linked to the PDMS show significant differences between cells seeded on treated FN regardless of rigidities (ZD, n>39). ***p<0.0001, **p<0.001, *p<0.05, n.s. = non-significant.

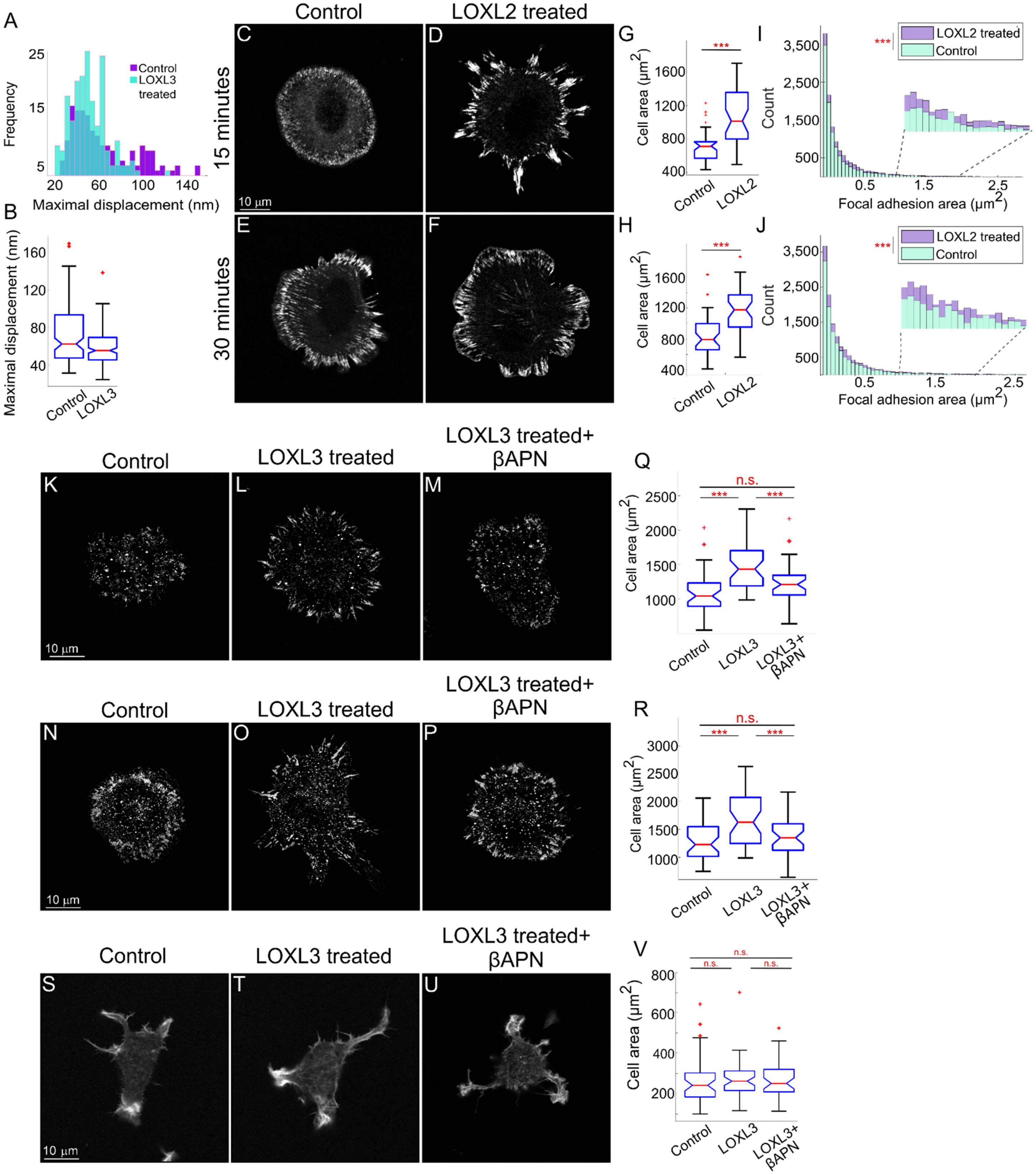

A
Control
LOXL3 treated
Frequency
Maximal displacement (nm)
B
Maximal displacement (nm)
Control
LOXL3
Control
LOXL2 treated
15 minutes
30 minutes
C
D
E
F
10 μm
G
H
Cell area (μm2)
LOXL2
I
J
LOXL2 treated
Control
Count
Focal adhesion area (μm2)
***
K
L
M
N
O
P
S
T
U
LOXL3 treated
LOXL3 treated+ βAPN
Q
R
V
n.s.
LOXL3+ βAPN

**Figure S2. Lysyl oxidases affect cell behavior specifically on FN coated substrates, Related to Figure 2.** Histogram and boxplot of the maximal displacement of pillars by cells seeded on LOXL3-treated laminin (A-B). Pax-GFP on sham- and LOXL2-treated FN, seeded on glass after 15 minutes (C-D) and 30 minutes (E-F). Quantifications of cell area show significant differences in cell area of those seeded on the treated FN (G-H, n=40). Histogram of focal adhesion area of cells seeded on sham- or LOXL2-treated FN demonstrate that LOXL2 treatment promotes larger adhesions (I-J). Paxillin immunostaining of wild-type MEFs (K-M) and human dermal fibroblasts (N-P) on sham- and LOXL3-treated FN with or without βAPN, seeded on glass and fixed after 15 minutes. Quantifications of cell area show significant differences in cell area of those seeded on the treated FN (Q-R, n>38). Pax-GFP on sham- and LOXL3-treated laminin with or without βAPN, seeded on glass and fixed after 15 minutes (S-U). Quantifications of cell area show no difference between cells seeded on treated laminin compared to controls (V, n>34). ***p<0.0001, n.s. = non-significant.

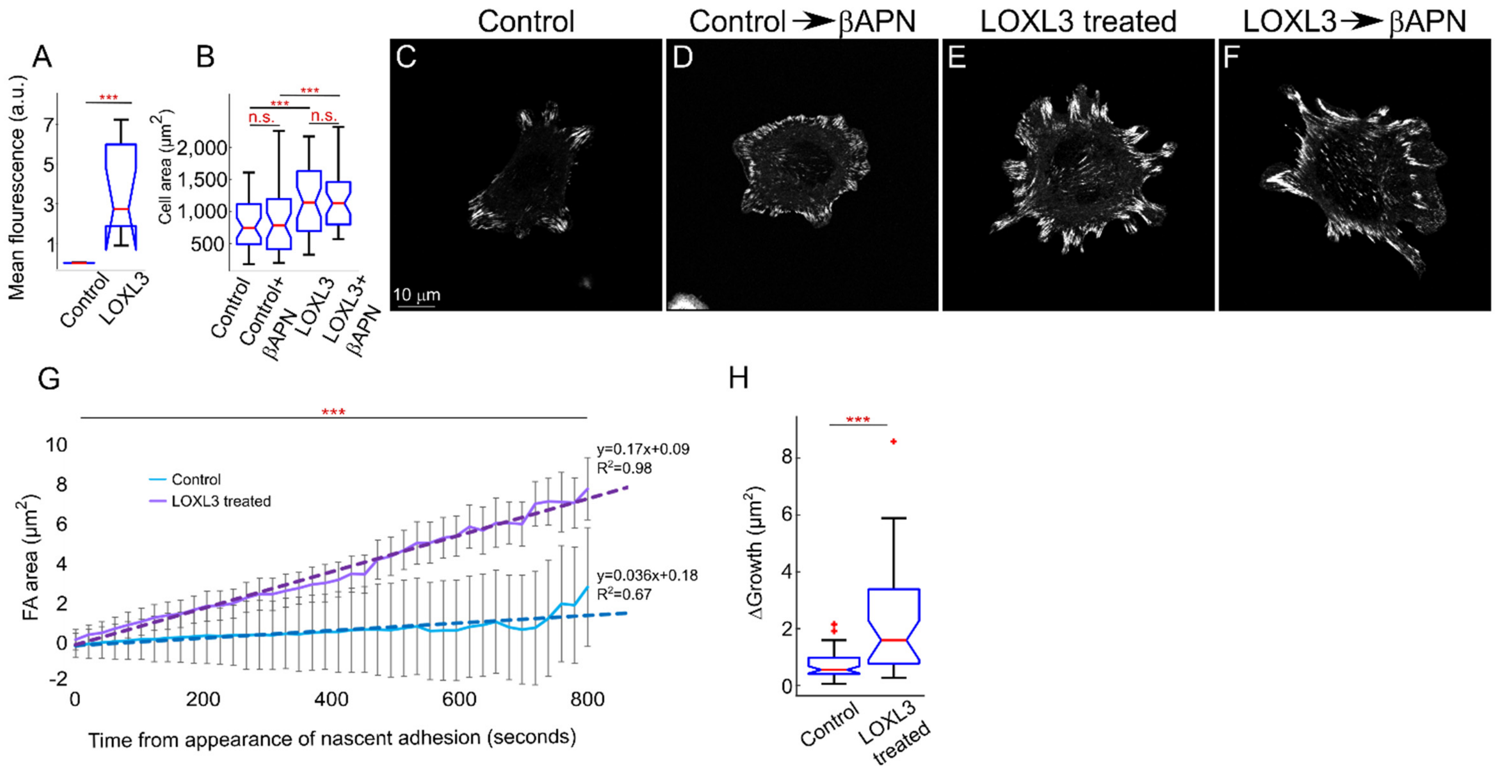


**Figure S3. Lysyl oxidases activity affects cell adhesion directly via changes in FN, Related to Figure 2.** Quantification of LOXL3 following treatment and washes (A, n=10). Quantification of cell area shows that no change was observed following late βAPN incubation, demonstrating that Lox activity regulates cell adhesion directly via FN rather than through integrin activation (B, n=40). Pax-GFP on sham- and LOXL3-treated FN followed by additional βAPN incubation on glass, fixed after 30 minutes (C-F). Graph shows confocal live-imaging quantifications of nascent adhesion area over time of cells seeded on FN with or without LOXL3 treatment, demonstrating the significant elevation in adhesion growth rate following LOXL3 treatment. Linear equations and R2 values were added to each treatment (G). Boxplot representing the delta growth (the difference between the maximal area and the minimal area) of nascent adhesions following live-imaging and adhesion analysis (H). ***p<0.0001, n.s. = non-significant.

A

**Untreated FN — Intensity values**

| Gene names | Position within protein | FN_1 | FN_2 | FN_3 |
|---|---|---|---|---|
| FN1 | 116 | 2.98E+05 | 1.63E+06 | 1.60E+07 |
| FN1 | 2391 | 7.82E+05 | 7.14E+06 | 1.62E+07 |
| FN1 | 2401 | 0.00E+00 | 7.14E+06 | 1.44E+07 |

**LOXL3 treated FN**

| Gene names | Position within protein | FN+ LOXL3_1 | FN+ LOXL3_2 | FN+ LOXL3_3 |
|---|---|---|---|---|
| FN1 | 116 | 4.11E+08 | 1.41E+06 | 1.94E+08 |
| FN1 | 2391 | 5.14E+07 | 8.24E+06 | 6.91E+07 |
| FN1 | 2401 | 7.10E+07 | 1.30E+07 | 8.33E+07 |

**Ratio mod/base**

| Position within protein | Average_ untreated | STD_ untreated | Average_ LOXL3 | STD_LOXL3 | Ratio LOXL3/ untreated |
|---|---|---|---|---|---|
| 116 | 1.78E-03 | 2.64E-03 | 2.42E-02 | 2.50E-02 | 13.61 |
| 2391 | 1.52E-02 | 1.35E-02 | 1.14E-01 | 1.48E-01 | 7.49 |
| 2401 | 9.03E-03 | 7.44E-03 | 5.23E-02 | 7.05E-02 | 5.79 |

**Untreated FN — Intensity values**

| Gene names | Position within protein | FN_1 | FN_2 | FN_3 |
|---|---|---|---|---|
| FN1 | 116 | 8.94E+06 | 6.92E+06 | 1.48E+07 |
| FN1 | 2391 | 1.89E+06 | 1.78E+06 | 2.83E+06 |
| FN1 | 2401 | 1.05E+07 | 1.06E+07 | 1.22E+07 |

**LOXL2 treated FN**

| Gene names | Position within protein | FN+ LOXL2_1 | FN+ LOXL2_2 | FN+ LOXL2_3 |
|---|---|---|---|---|
| FN1 | 116 | 4.11E+07 | 2.37E+07 | 4.43E+07 |
| FN1 | 2391 | 4.17E+07 | 1.40E+07 | 3.57E+06 |
| FN1 | 2401 | 5.03E+07 | 1.79E+07 | 1.25E+07 |

**Ratio mod/base**

| Position within protein | Average_ untreated | STD_ untreated | Average_ LOXL2 | STD_ LOXL2 | Ratio LOXL2/ untreated |
|---|---|---|---|---|---|
| 116 | 1.02E+07 | 4.11E+06 | 3.64E+07 | 1.11E+07 | 3.56 |
| 2391 | 2.16E+06 | 5.79E+05 | 1.98E+07 | 1.97E+07 | 9.14 |
| 2401 | 1.11E+07 | 9.74E+05 | 2.69E+07 | 2.05E+07 | 2.42 |

B

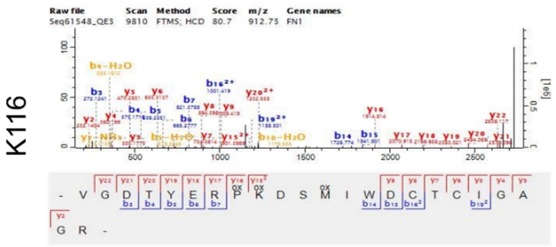


C

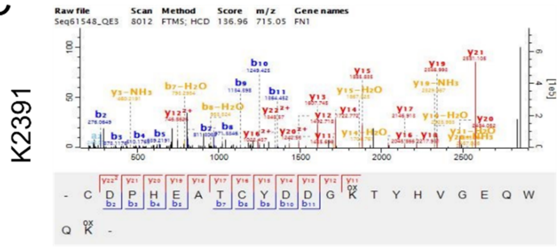


D

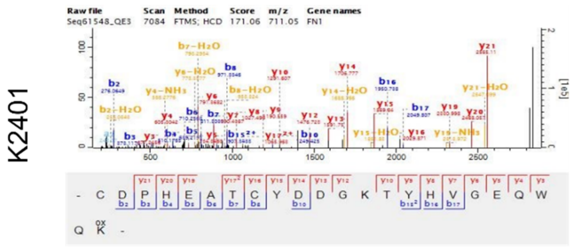


E

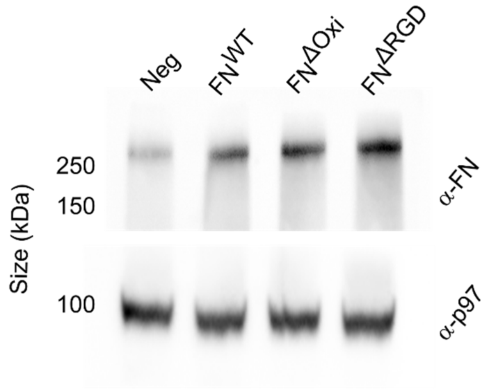


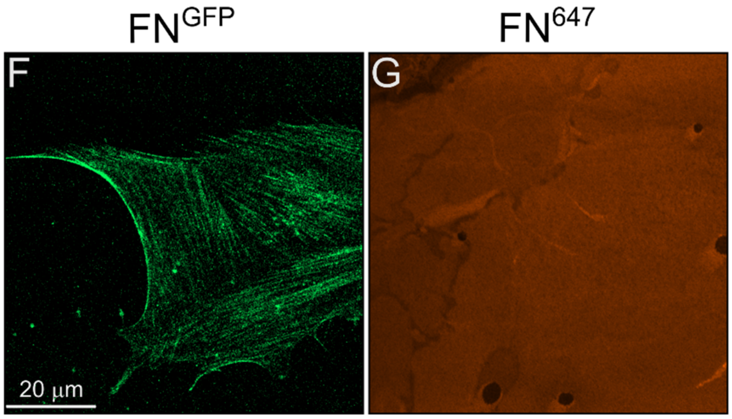

**Figure S4. Lysyl oxidases specifically oxidize FN, Related to Figure 3.** Representative intensity values of oxidized peptide identification in untreated (top) and treated (middle) FN. The tables on the left refer to experiments that involved LOXL3 and those on the right refer to experiments that involved LOXL2. The bottom tables show the averages and standard deviations (std) of the modified (mod, oxidized) to non-modified (base, non-oxidized) ratios in the samples (A). Representative MS/MS spectra of identified peptides (B-D). FN-GFP cells seeded on FN-647 and fixed after 48 hours (E-F).

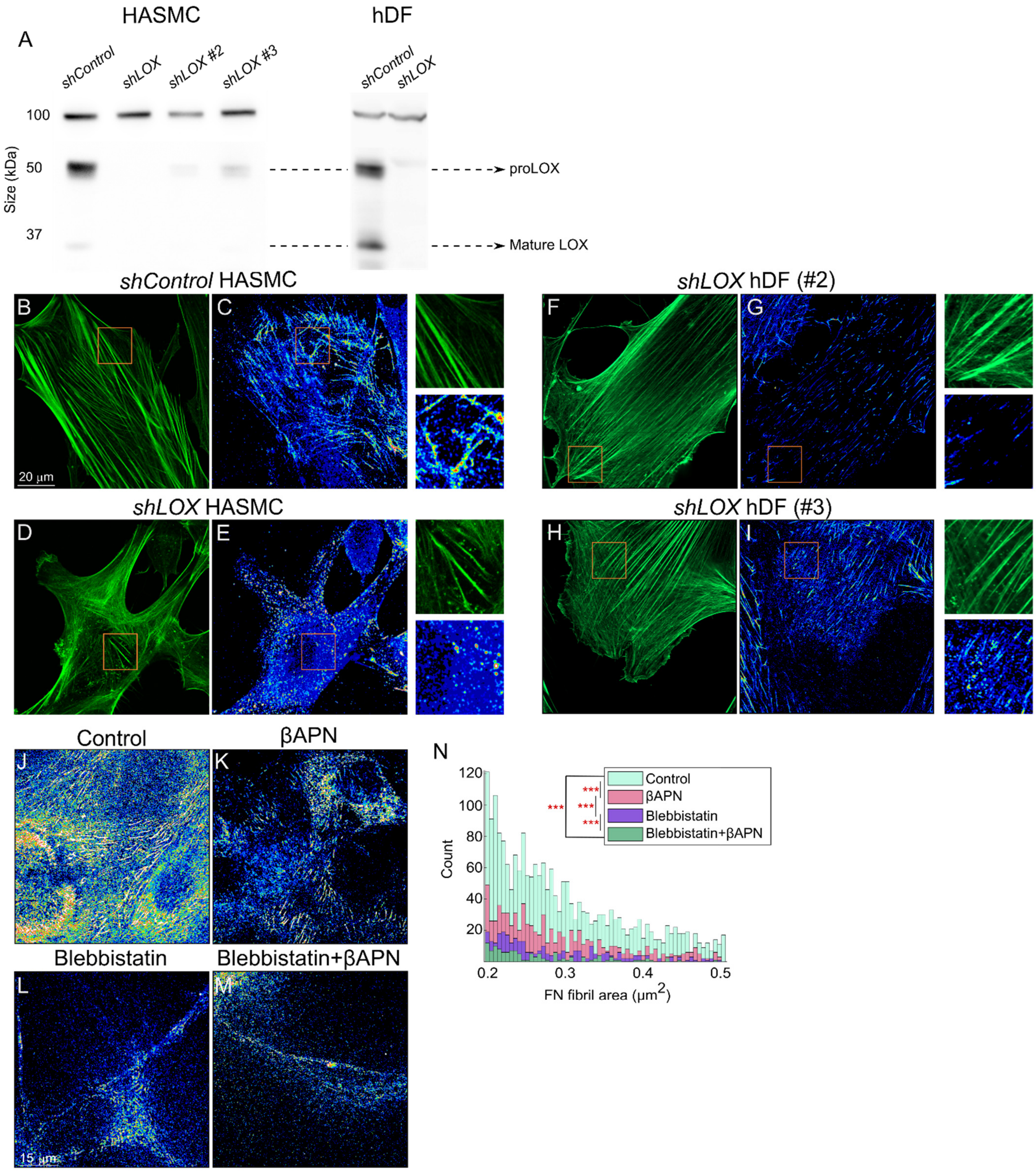
HASMC
hDF
A
shControl
shLOX
shLOX #2
shLOX #3
shControl
shLOX
100
50
37
Size (kDa)
proLOX
Mature LOX
shControl HASMC
B
C
20 μm
shLOX hDF (#2)
F
G
shLOX HASMC
D
E
shLOX hDF (#3)
H
I
Control
βAPN
J
K
Blebbistatin
Blebbistatin+βAPN
L
M
15 μm
N
Control
βAPN
Blebbistatin
Blebbistatin+βAPN
***
***
***
***
Count
120
100
80
60
40
20
0.2
0.3
0.4
0.5
FN fibril area (μm2)

**Figure S5. Lysyl oxidases are essential for long-term FN fibrillogenesis, Related to Figure 4.** Western blot against LOX from lysates of human dermal fibroblasts (hDF) expressing *shControl* and each of the three different *shLOX* lentiviral plasmids (A). Actin and FN immunostaining (color-coded for intensity) of shControl (B-C) and shLOX (D-E) HASMC seeded on glass without external FN coating and fixed after 24 hours. Images on the right are zoom-ins of the orange boxes in each image. Actin and FN immunostaining (color-coded for intensity) of *shLOX* #2 (F-G) and *shLOX* #3 (H-I) hDF seeded on glass without external FN coating and fixed after 24 hours. Images on the right are zoom-ins of the orange boxes in each image. Pax-GFP cells treated with blebbistatin, βAPN, or both, seeded on glass and fixed after 24 hours (F-I). Histogram of the area of FN fibrils generated by Pax-GFP MEF treated with blebbistatin, βAPN, or both, showing the significant reduction of fibril formation with the combination of both treatments (J). ***$p<0.0001$, n.s. = non-significant.
